# Endothelial Nitric Oxide Synthase (eNOS) S1176 phosphorylation status governs atherosclerotic lesion formation

**DOI:** 10.1101/2022.01.13.476234

**Authors:** Tung D. Nguyen, Nur-Taz Rahman, William C. Sessa, Monica Y. Lee

**Author notes:** Corresponding author: Monica Y. Lee, Ph.D., Center for Cardiovascular Research, University of Illinois at Chicago, College of Medicine Research Building 835 S. Wolcott Ave., Chicago, IL 60612.

## Abstract

**Objective:** We have previously demonstrated the *in vivo* importance of the Akt-eNOS substrate-kinase relationship, as defective postnatal angiogenesis characteristic of global Akt1-null mice is rescued when bred to ‘gain-of-function’ eNOS S1176D mutant mice. While multiple studies support the vascular protective role of endothelial NO generation, the causal role of Akt1-dependent eNOS S1176 phosphorylation during atherosclerotic plaque formation is not yet clear.

**Approach & Results:** We herein bred congenic ‘loss-of-function’ eNOS S1176A and ‘gain-of-function’ eNOS S1176D mutant mice to the exacerbated atherogenic Akt1^-/-^; ApoE^-/-^ double knockout mice to definitively test the importance of Akt-mediated eNOS S1176 phosphorylation during atherogenesis. We find that a single amino acid substitution at the eNOS S1176 phosphorylation site yields divergent effects on atherosclerotic plaque formation, as an eNOS phospho-mimic aspartate (D) substitution at S1176 leads to favorable lipid profiles and decreased indices of atherosclerosis, even when on a proatherogenic Akt1 global deletion background. Conversely, mice harboring an unphosphorylatable mutation to alanine (S1176A) result in increased plasma lipids, increased lesion formation and cellular apoptosis, phenocopying the physiological consequence of eNOS deletion and/or impaired enzyme function. Furthermore, gene expression analyses of whole aortas indicate a combinatorial detriment from NO deficiency and Western Diet challenge, as ‘loss-of-function’ eNOS SA mice on a Western diet present a unique expression pattern indicative of augmented T-cell activity when compared to eNOS S1176D mice.

**Conclusions:** By using genetic epistasis approaches, we conclusively demonstrate that Akt-mediated eNOS S1176 phosphorylation and subsequent eNOS activation remains to be the most physiologically relevant method of NO production to promote athero-protective effects.

## INTRODUCTION

Endothelial cells (EC) play a vital role in modulating vascular responses. The importance of endothelial nitric oxide (NO) production in cardiovascular protection has been long-established through investigation of numerous physiological models. EC-derived NO is critical for the regulation of several vascular responses, including vascular tone, blood flow, leukocyte-endothelial interactions, platelet adhesion/aggregation, and vascular smooth muscle cell function. A decrease in NO bioavailability results in endothelial dysfunction, as characterized by features conducive to the development of atherosclerosis to influence parameters such as thrombosis, inflammation, neointimal proliferation, and vasoconstriction^1,2^.

eNOS knockout mice demonstrate the most extreme outcomes of eNOS loss, such as increased leukocyte-endothelial interaction, hypertension and atherosclerosis^3,4^. Overexpression of eNOS also, paradoxically, yields larger atherosclerotic lesions, which was later attributed to consequential eNOS dysfunction and uncoupling^5^. eNOS uncoupling is therefore an important mechanism underlying the pathogenesis of EC dysfunction and downstream atherogenesis, where increasing evidence suggests that proper functionality of the eNOS enzyme, rather than expression levels, is most critical for cardiovascular homeostasis^6^. Among the various mechanisms of regulation, eNOS protein phosphorylation at serine 1176 (1179 in bovine, 1177 in human and 1176 in murine) plays a critical role in eNOS enzymatic activity^7,8^. Various extracellular stimuli (*e.g.* insulin, VEGF, shear stress) generate NO through eNOS S1176 phosphorylation, where several kinases (*e.g.* Akt, PKA, PKG, AMPK, CaMKII, etc.) have been identified, implying the physiological importance of eNOS phosphorylation and activation^9^.

Although several kinases can phosphorylate eNOS at S1176, our previous studies using genetic epistasis approaches demonstrate that the kinase Akt is the most critical for eNOS phosphorylation and vascular function *in vivo*. We have shown that the impaired angiogenic phenotypes typically seen in Akt1 deficient mice are rescued when crossed with eNOS ‘gain-of-function’ (serine to aspartate, S1176D) mice^8^. The observed rescue in impaired angiogenesis demonstrates the importance of Akt1 activity as the predominant kinase for eNOS phosphorylation in vascular repair. While the Akt-eNOS relationship has been described in the context of adaptive angiogenesis, the causal role of Akt-dependent eNOS phosphorylation in atherogenesis has not yet been investigated using these available mouse models. The definitive role of the Akt1 signaling pathway in atherosclerosis remains uncertain, as several *in vitro* studies suggest Akt may serve either an athero-protective or athero-prone role^10–12^. Moreover, the conflicting results using eNOS overexpression systems emphasizes the importance of proper eNOS function, rather than overt modulation of protein levels^3,6^.

In this study, we aimed to investigate the importance of the Akt1-eNOS activation cascade on atherosclerotic plaque development using genetically modified mice. Knock-in mice carrying the described ‘gain-of-function’ (S1176D) or a ‘loss-of-function’ (S1176A) mutations in the endogenous eNOS gene were bred to the previously published Akt1^-/-^; ApoE^-/-^ double knockout mice^13^, a model of severe atherosclerosis. These triple transgenic mice were developed to address the importance of the Akt1-eNOS activation cascade in an extreme model of murine atherosclerosis using genetic means. Use of an Akt1-null background will also eliminate other means of Akt-mediated vascular protection to fully define the role of eNOS S1176 phosphorylation in atherogenesis. We herein report that a single amino acid substitution at the eNOS S1176 phosphorylation site yields divergent effects on plasma lipids and atherosclerotic plaque formation when placed on an Akt1^-/-^; ApoE^-/-^ double knockout genetic background. Our study provides the first *in vivo* evidence that the preservation of eNOS function is essential to mitigate atherosclerotic lesion formation, despite the global loss of Akt1 expression. Moreover, gene expression analyses indicate that impaired eNOS phosphorylation together with a Western Diet challenge promotes vascular signatures reflective of the adaptive immune response. We herein demonstrate that Akt1-directed eNOS activation indeed serves an athero-protective role, where we further substantiate the Akt1-eNOS axis as the major signaling mechanism that links endothelial integrity to cardiovascular disease outcome.

## MATERIALS & METHODS

All gene expression data will be made publicly available at GEO.

The data that support the findings of this study are available from the corresponding author upon request.

### Animal Procedures

Mice expressing an endogenous phosphomimetic (S1176D) and unphosphorylatable (SA) eNOS point-mutations at the S1176 site were backcrossed to a C57Bl/6J background (provided by Paul L. Huang^8,14^). Akt1^-/-^; ApoE^-/-^ double knockout mice were previously backcrossed onto a C57Bl/6J background and utilized for the studies herein^13^. These triple allele mice hence reflect eight generations of backcrossing to a C57Bl/6 background. Published studies report a significant decrease in litter size in both the eNOS^-/-^ and global Akt1^-/-^ mice^15–18^. Akt1 heterozygous females (SA [or S1176D]^+/+^; Akt1^+/-^; ApoE^-/-^) were therefore bred to Akt1 homozygote KO males (SA [or S1176D]^+/+^; Akt1^-/-^; ApoE^-/-^) to maximize the chances of viable offspring. Although genetic crossing strategies were employed, the phospho-impaired eNOS S1176A mutation on a sub-optimal Akt1 expression system resulted in significantly decreased litter and offspring numbers (average n=3 pups per litter, *unpublished*). Furthermore, our atherosclerosis studies required use of male offspring harboring homozygote triple-allelic mutations (SA [or SD]^+/+^; Akt1^-/-^; ApoE^-/-^) to minimize variability and additionally maintain the difficult genetic lines, hence the low numbers in these studies.

Triple allelic homozygote mice were maintained on standard laboratory diet (Teklad 7912, Envigo), where at ∼8wks of age mice were fed *ad libitum* with a high-fat (40% kcal), high-cholesterol (1.25%) Western Diet (Research Diets, D12108). Adult mice were fed with Western Diet for ∼4 or 12 weeks for atherosclerosis studies. All experiments were approved by the Institutional Animal Care Use Committee at Yale University.

### Blood plasma measurements

Mice were fasted for ∼12 hours prior to blood collection. Blood samples were collected via retro-orbital bleeding on anesthetized mice. Samples (∼200uL/mouse) were transferred to Eppendorf tubes containing ∼2uL 0.5M EDTA, gently mixed, and immediately placed on ice. Blood samples were centrifuged for 15 minutes at 2000g at 4C, and plasma supernatant fractions were collected for analyses. Plasma NO levels were assessed using the Nitrate/Nitrite Colorimetric Assay Kit according to the manufacturer’s instructions (Caymen, 780001). Plasma triglyceride levels were determined using the Triglyceride (TG) Colorimetric Assay Kit (Caymen, 10010303) by measuring absorbance at 530nm and comparing values to TG standards. Plasma cholesterol levels were similarly determined using the Total Cholesterol E Assay Kit (Fujifilm, 999-02601) by measuring absorbance at 600nm and comparing values to cholesterol standards.

### Oil Red O en face staining

After a 12-week Western Diet challenge, mice were anesthetized (ket/xyl) and perfused with PBS (pH 7.4) prior to removal of the aorta. Aortas were dissected from the aortic valve to the iliac bifurcation and fixed in 4% PFA/PBS (pH 7.4) overnight at 4°C. Aorta samples were thoroughly cleaned prior to longitudinal opening and pinning down in silicone plates for *en face* preparation. Lipid-rich lesions were identified using previously described Oil Red O staining techniques^13^. Images were acquired using a Nikon SMZ 1000 microscope and lesion areas were blindly quantified using ImageJ software. Lesion areas are reported as percentages of total aortic area. Data reflect multiple cohorts of mice.

### Histological analysis

The aortic root and brachiocephalic arteries were isolated after a 12-week Western Diet challenge, similar to aorta harvest. After fixation, tissues were incubated overnight at 4C in 30% sucrose/PBS and embedded in OCT medium for storage at -80C. Tissues were sectioned (6μm) and analyzed for plaque burden via immunohistochemistry at 3 different equally spaced intervals (∼120μm), similar to previous methods^19^. Total lesion area (from internal elastic lamina to the lumen) and acellular/necrotic areas were measured by hematoxylin-eosin (H&E) and Masson’s Trichrome staining, as previously described^20^. In brief, the necrotic cores were defined as a clear area devoid of staining, where outlines were determined by implementing a 3,000 μm^2^ threshold to avoid areas that likely do not represent necrotic core regions. Total lesion and necrotic core areas were measured using ImageJ.

### Immunostaining

Mice were anesthetized (ket/xyl) and perfused with PBS (pH 7.4) prior to removal of target organs (*e.g.* brachiocephalic artery, heart). Samples were fixed in 4% PFA/PBS overnight at 4°C followed by immediate dehydration in 30% sucrose/PBS for OCT embedding. Frozen tissues were cut at 5um sections and stained with hematoxylin/eosin or Masson’s Trichrome. Adjacent sections were immunostained with the following primary antibodies (4°C, overnight): anti-CD68 (Serotec, MCA1957); anti-VCAM1 (BD 550547); anti-ITGA5 (BD 555615); anti-Stat3 (Cell Signaling 12640); anti-RelA (Proteintech 10745-1-AP); anti-Mac2 (Cedarlane CL8942AP). Tissue sections were subsequently incubated with corresponding AlexaFluor secondary antibodies (2hrs, room temp). Cell apoptosis was measured using an *in situ* cell death detection kit and following the manufacturer protocol (Roche, 12156792910). Images were obtained using an inverted fluorescent microscope (Leica DMi8) and quantified via ImageJ. For all parameters, multiple sections were assessed for each tissue sample where the average values are reported for each mouse. Data reflect multiple cohorts of mice.

### Gene Expression Analyses

Upon euthanasia (ket/xyl), mice were perfused with PBS and aortic tissue (ascending aorta to the femoral arteries) was quickly isolated. Whole aorta samples were thoroughly cleaned of perivascular fat and immediately flash-frozen in liquid nitrogen to minimize technical effects imposed by time (within 20 minutes of euthanasia). Whole aorta tissues were then processed for RNA extraction and quality checks. RNA quality was assessed using spectrophotometric (NanoDrop) readings to ensure that all samples were at A260/280 ratios of ∼2.0. Satisfactory samples were prepared for hybridization reactions with the NanoString platform (PanCancer Immune Profile Mouse). In brief, the NanoString nCounter Gene Expression panels allow for the analyses of up to ∼800 RNA targets selected for published significance in key biological pathways using a digital detection and direct molecular barcoding approach. Normalized expression counts were extracted from the NanoString nSolver software for further data analyses using Qlucore Omics Explorer® (Version 3.8). Samples were log2 transformed in Qlucore and assigned into their respective groups: SA+Standard (2 samples), SD+Standard (3 samples), SA+WD (3 samples), and SD+WD (3 samples). The 783 genes in the dataset were used to generate PCA plots and heatmaps. Best clustering of the samples was achieved with multi-group (ANOVA) comparison of all 4 groups at a p-value ≤ 0.05 (corresponding q-value ≤ 0.125), which identified 305 differentially expressed genes. PCA plots are shown in **Supplemental Figure 1**. Sample groups were further analyzed using two group comparisons to identify differentially expressed genes based on genotype or diet conditions. Differentially expressed genes were then subjected to Venn diagram analyses to identify relationships amongst the sample conditions. Heatmaps were generated for select overlapping gene lists across all sample conditions, where hierarchical clustering was applied for the identified genes. Post hoc Tukey HSD analysis was applied to test significance between groups. Pathway analyses were carried out for these different gene sets using Ingenuity Pathway Analysis (IPA, Qiagen, MD). From IPA we obtained predicted canonical pathways and upstream regulators.

### Statistical Anaylsis

All data are shown as mean ± standard error of the mean (SEM) using GraphPad Prism software (Version 9.5, GraphPad). Statistical significance was evaluated using an unpaired two-tailed Student’s t test for two-group comparisons of normally distributed data with equal variance. Statistical significance was evaluated using a 2-Way ANOVA followed by Bonferroni’s post-test for multi-group comparisons with equal variances. Significance was determined based on a p value calculation of p<0.05.

## RESULTS

### eNOS-S1176D ‘gain-of-function’ mutant mice show favorable plasma profiles

Mice expressing an endogenous phosphomimetic (S1176D) and unphosphorylatable (S1176A) eNOS point-mutation at S1176 were crossed with Akt1^-/-^; ApoE^-/-^ double knockout mice to generate triple allelic homozygote mice. Breeding methods were taken to maximize experimental mouse numbers (see Methods). With these newly generated mice, this study aims to clarify the importance of eNOS as an Akt1 substrate by using a previously published Akt1-deficient atherosclerotic background (i.e. Akt1^-/-^; ApoE^-/-^ double knockout)^13^. More importantly, these triple allelic homozygote mice carry knock-in mutations of the critical Akt1 phosphorylation site on eNOS (S1176) to render the enzyme ‘constitutively active’ or ‘less active’. Male mice at ∼8 weeks of age were fed a Western Diet for a duration of 12 weeks. A 12-week Western Diet leads to a significant increase in body weight in both eNOS mutant groups when compared to pre-Western Diet conditions. Manipulation of eNOS, however, had no effect on total weight, irrespective of the diet type (**Figure 1A**). We also did not observe any significant mortality at the end of the 12-week feeding period. Introduction of these eNOS mutations do not affect the levels of eNOS protein levels, but rather modulates eNOS activity through manipulation of the S1176 site, as previously shown^8^. To confirm the enzymatic status of our eNOS mutant mice triple allelic mice, blood plasma was collected, where results indicate significantly higher NO levels in S1176D mice when compared to S1176A mice both pre-and post-Western Diet fed conditions **(Figure 1B)**. A 12-week Western Diet challenge, however, leads to a significant reduction in plasma NO levels in eNOS S1176A ‘loss-of-function’ mice when compared to standard diet conditions. This is not observed in the eNOS S1176D ‘gain-of-function’ mice. eNOS S1176D mice also exhibit a trending decrease in plasma cholesterol levels and significantly less triglycerides under standard diet conditions **(Figure 1C&1D)**. A 12-week Western Diet challenge leads to a significant increase in plasma cholesterol in both eNOS mutant groups and significantly increases plasma triglycerides only in eNOS S1176A mutant mice. When comparing eNOS mutant groups under Western Diet fed conditions, total plasma cholesterol and triglycerides are significantly lower in eNOS S1176D mice **(Figure 1C&1D)**. These results demonstrate the importance of eNOS S1176-mediated enzyme activation on generating plasma NO and limiting plasma atherosclerotic lipid mediators.

**Figure 1.**
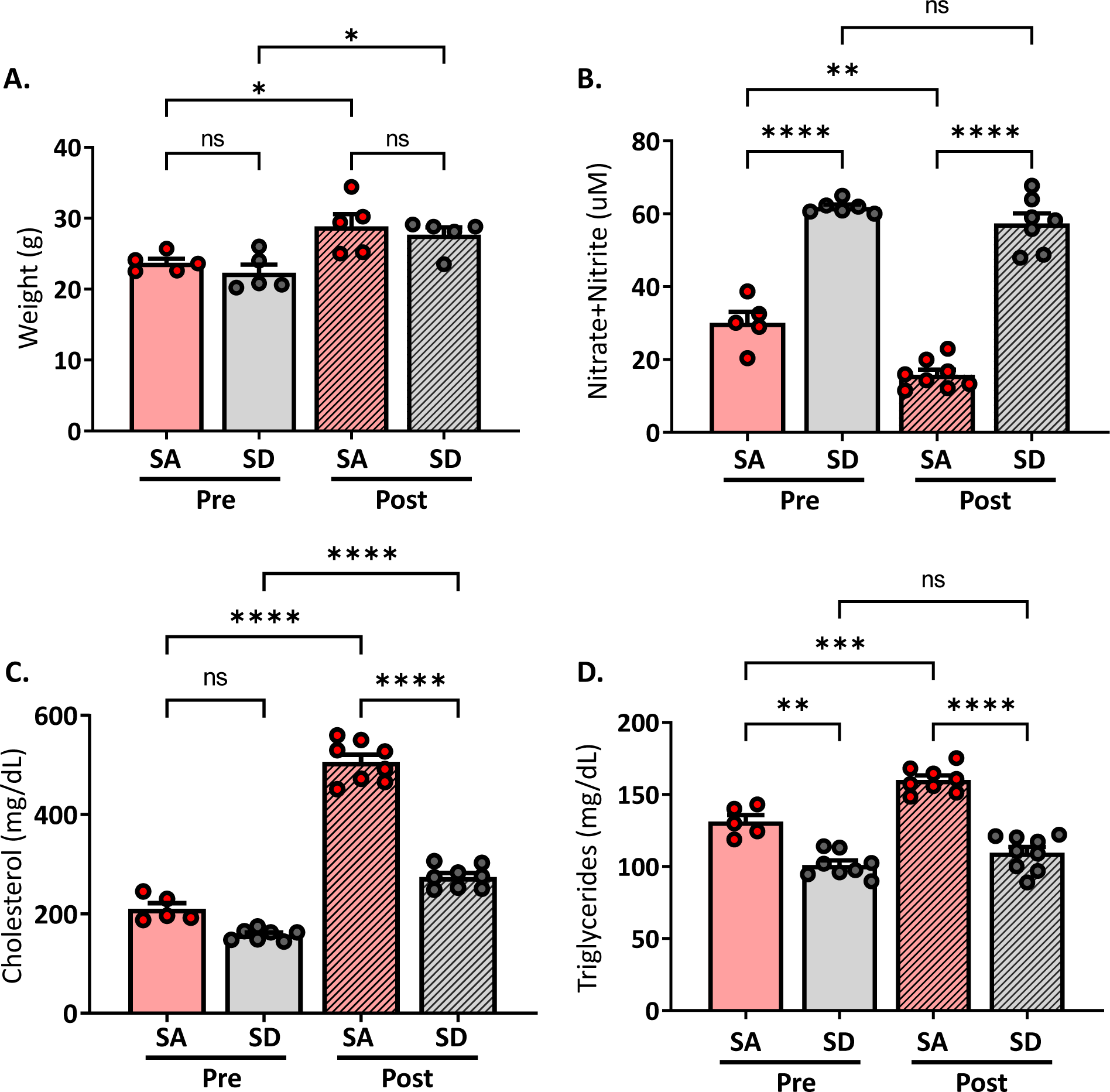
Weights and plasma measurements in eNOS mutant mice. **(A)** Recorded mouse body weights pre- and post-12 week Western Diet feeding. Plasma measurements of **(B)** nitric oxide show significantly higher levels in eNOS S1176D gain-of-function mice with both chow and Western Diet fed conditions. **(C)** Total plasma cholesterol **(D)** and triglyceride levels in mice pre- and post-12 week Western Diet feeding. *n=5 to 8 mice per group, **** p<0.0001, *** p<0.001, ** p<0.01*

### eNOS-S1176A phospho-impaired mutant mice display increased aortic lesion formation

Aortas were isolated after a 12-week Western Diet feed period to visually inspect atherosclerotic lesion formation in aortic regions and areas of lesion predilection, namely sites of branching and vessel curvature. *En face* Oil Red O staining of aortas from eNOS S1176A mice show larger areas of aortic lesions when compared to eNOS S1176D mice (**Figure 2A&2B**). Lesion quantifications show significantly increased plaque areas in eNOS S1176A mice throughout regions of the aortic arch (**Figure 2C**), thoracic aorta (**Figure 2D**), and abdominal aorta (**Figure 2E**). Additional whole aorta images from experimental groups can be found in **Supplemental Figure 2**. Cross-sectional analyses of the aortic sinus indicate that ‘gain-of-function’ eNOS S1176D mice develop visibly decreased atherosclerotic lesions and lipid deposition when compared to eNOS S1176A mice (**Figures 2F&2G**), corroborating the decreased plaque formation seen throughout the whole aorta.

**Figure 2.**
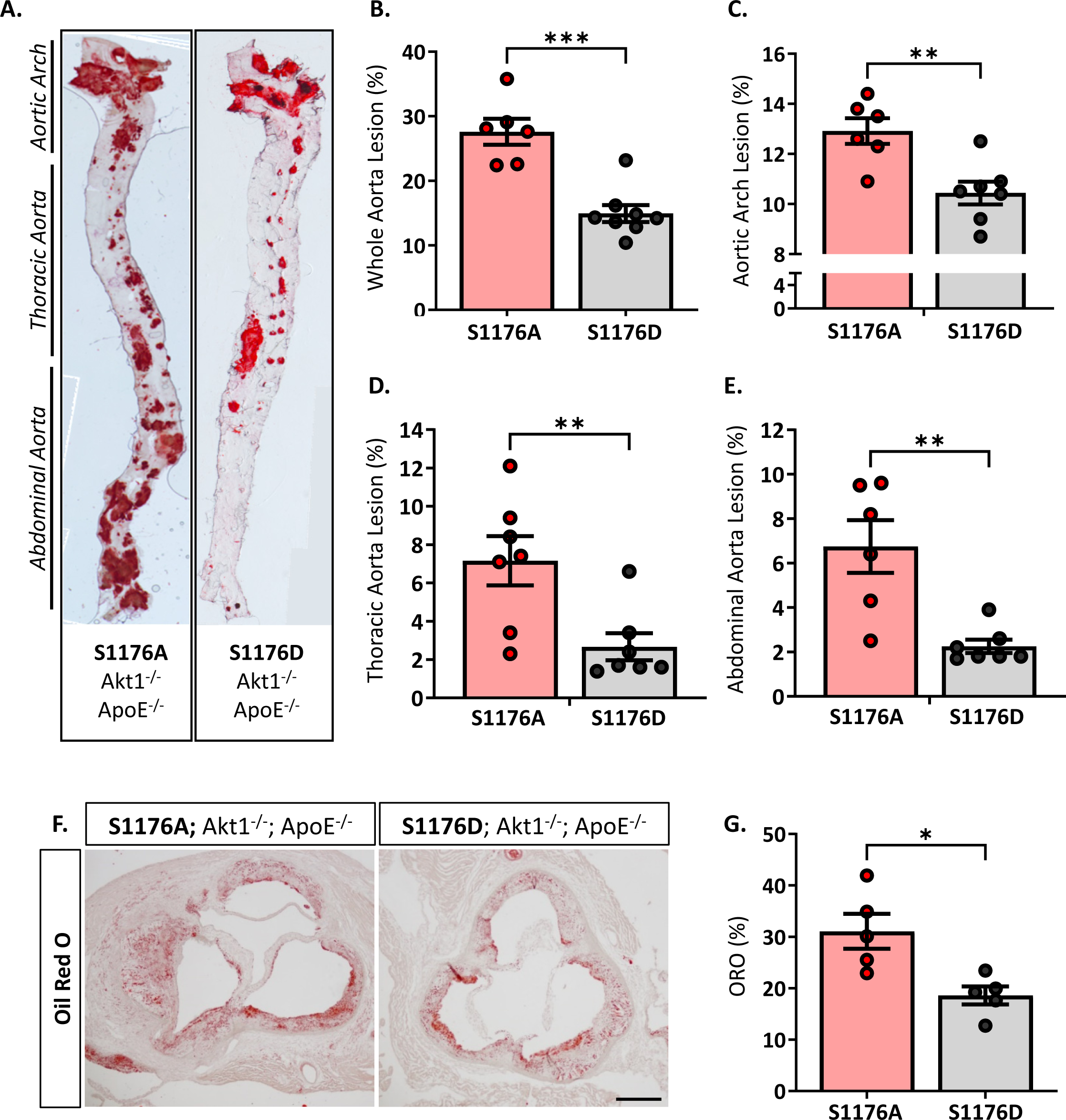
Increased lesion formation and lipid deposition in eNOS-S1176A phospho-impaired mutant mice. **(A)** *En face* staining for Oil Red O in aortas from eNOS mutant mice indicate significant lesion formation throughout the aorta in S1176A compared to S1176D mutant mice. Quantified in **(B-E). (F)** Oil Red O staining of the aortic root 12 weeks post-Western Diet challenge. Quantified in **(G**). *n=6-8 per group, *** p<0.001, ** p<0.01, *p<0.05*

### Atherosclerotic mice expressing the ‘gain-of-function’ eNOS mutation (S1176D) exhibit less plaque necrosis apoptosis and inflammation

Expression of the ‘gain-of-function’ eNOS S1176D mutant results in decreased necrotic core regions, as shown by Masson’s Trichrome staining (**Figures 3A-C**). Atherosclerotic necrotic core regions were defined as a clear area devoid of staining, as previously described^20^. We additionally examined cellular apoptosis using terminal deoxynucelotidyl transferase dUTP end-labeling staining techniques in adjacent cross sections of the aortic sinus, where eNOS S1176A mutant mice exhibit a significant increase in overall TUNEL+ plaque regions (**Figures 3D & 3E**). Cross-sectional analyses of the brachiocephalic artery also indicate significantly larger plaques in the eNOS S1176A mice along with increased lipid deposition and apoptosis (**Supp. Fig. 3**). Moreover, eNOS S1176A mice exhibit significantly higher levels of integrin-α5 (ITGA5), a fibronectin receptor associated with increased inflammation and atherosclerosis^21^ (**Figure 3F&3G**). These results suggest that impairment of the Akt-eNOS signaling axis in the triple allelic eNOS S1176A ‘loss-of-function’ mice contributes to exacerbated plaque formation.

**Figure 3.**
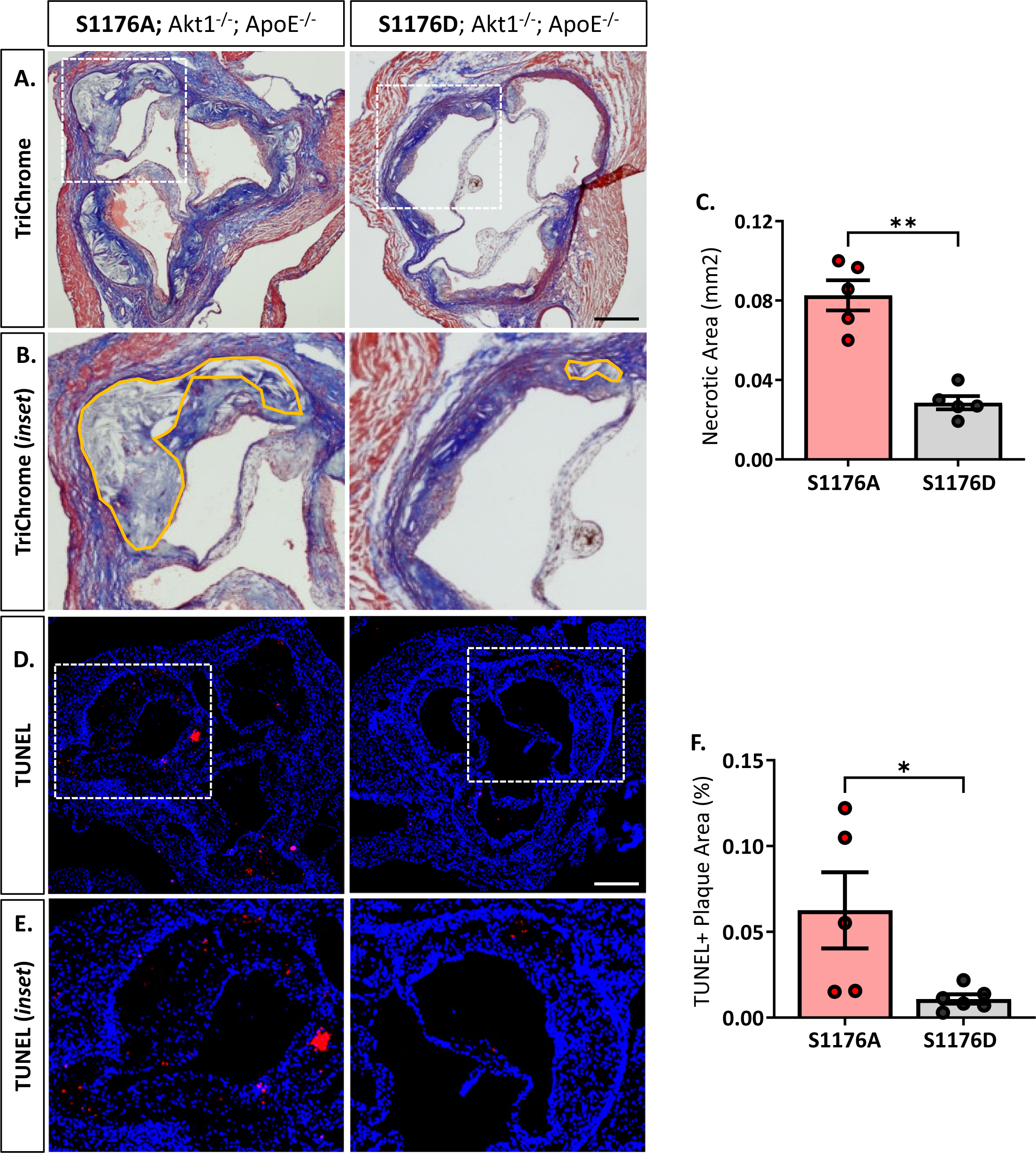
Increased necrotic core formation and cellular apoptosis in phospho-impaired eNOS-S1176A triple allelic mutant mice. **(A)** Trichrome staining of the aortic root 12 weeks post-Western Diet with **(B**) inset images of necrotic core regions drawn in orange. Quantified in **(C). (D)** TUNEL staining of the aortic root 12-weeks post-Western Diet, quantified in **(E). (F)** ITGA5 immunostaining of the brachiocephalic artery 12 weeks post-Western Diet feed indicates a significant increase in eNOS S1176A compared to S1176D mice. Quantified in **(G)**. *n = 5-6 per group, ** p<0.01, *p<0.05*

Next, we examined the expression of macrophage-like markers and VCAM-1 levels in the two strains. The greater lesion size in eNOS S1176A mice is accompanied by enhanced CD68 positive cells in the plaque and VCAM-1 levels, where these effects are reduced in eNOS S1176D mice (**Figure 4A-D**). eNOS S1176A mice also display a trending increase in Mac2 expression, suggesting an increase in monocyte infiltration (**Supp. Fig. 4**). The increase in inflammatory cell and VCAM1 expression corroborates previously reported increases of VCAM1 in Akt1-null mice, where enhanced proinflammatory gene expression was found to be secondary to macrophage infiltration^13^. We herein show that expression of the ‘gain-of-function’ eNOS S1176D mutation exhibits decreased indices of inflammation, despite the lack of Akt1 expression and the chronic environmental stress of a Western Diet challenge. Our findings support the long-held notion that increasing NO bioavailability via eNOS S1176 activation mitigates plaque formation, emphasizing the athero-protective role of eNOS function.

**Figure 4.**
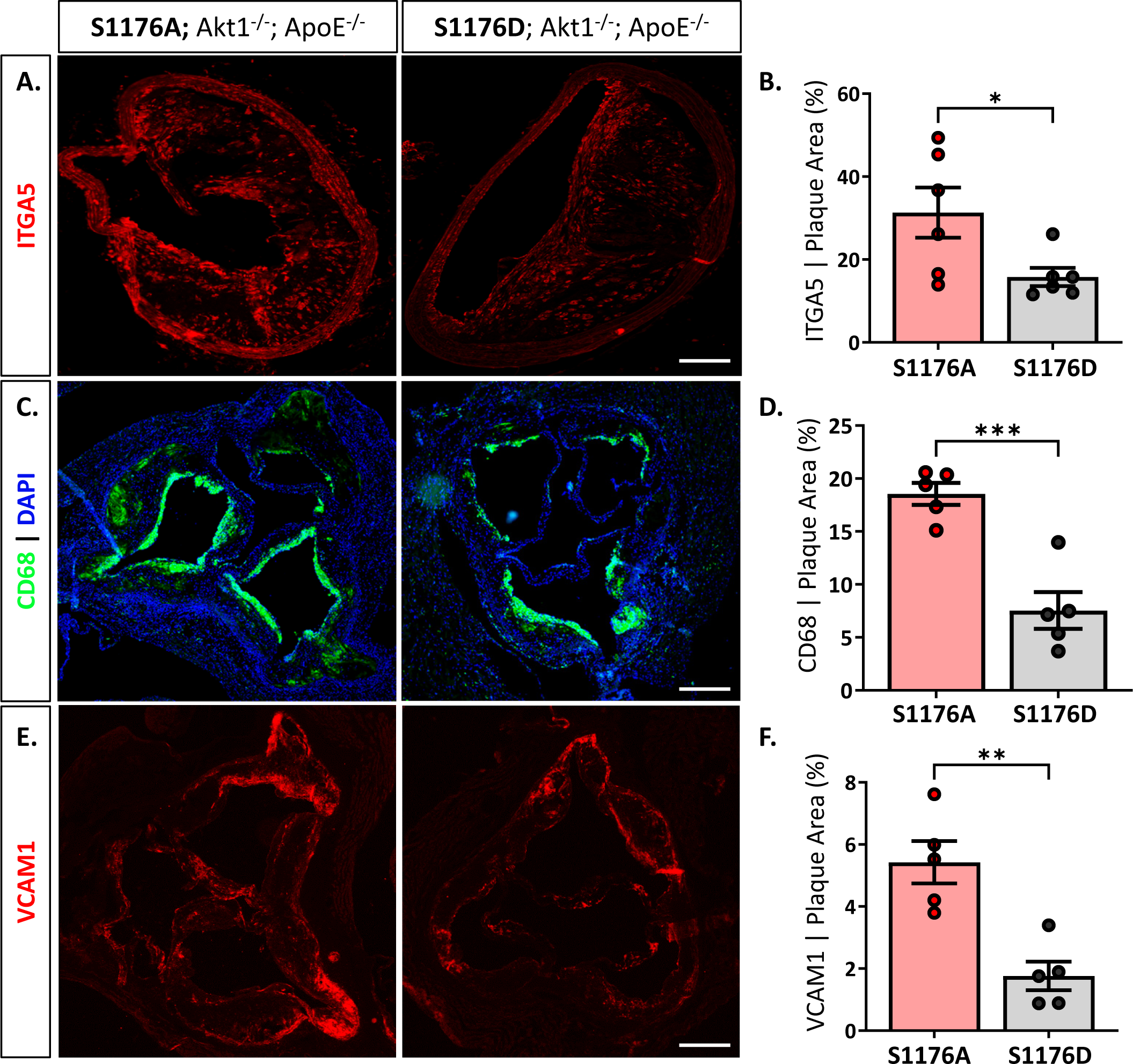
Increased CD68 and VCAM1 expression in phospho-deficient eNOS-S1176A mutant mice. Immunostaining of the aortic root shows enhanced expression of **(A)** CD68 and **(B)** VCAM1 within atherosclerotic lesions in eNOS S1176A mice. Quantified in **(C)** and **(D)**. *n = 5-6 per group, *** p<0.001, ** p<0.01*

Previous studies have shown that endothelial adhesion molecule expression and subsequent monocyte recruitment and proliferation requires at least 2 weeks of a hypercholesterolemic environment^22^. Moreover, adherent monocytes are found on the surface of activated endothelium in the thoracic aorta prior to the development of foam cell-rich fatty streaks within 4-weeks of Western Diet challenge in ApoE mice^23^. We therefore shortened the Western Diet feed duration to a 4-week duration to reflect early endothelial activation, and comprehensively profiled the transcriptional changes in whole aorta samples between comparison groups to capture the various mechanisms of atherogenesis. Importantly, we also isolated the whole aorta to account for the long-range vascular protective effects of eNOS activity and gaseous NO on the various cells of the vasculature. Samples were then processed on a NanoString Immune Profile platform to determine how eNOS dysfunction together with a 4-week Western Diet challenge affects aortic gene expression, with focus on adaptive and innate immune response genes.

To find discriminating variables, multigroup (ANOVA) workflow was applied across the four sample groups. A heatmap of the normalized expression data results in clustering of sample groups between the comparison groups, indicating sample similarity amongst biological replicates (**Figure 5A**, p≤0.05, 305 genes). More importantly, unsupervised hierarchical gene clustering highlights several unique gene clusters across the groups of interest. Gene expression counts were further analyzed using the Ingenuity Pathway Analysis (IPA) software to identify canonical pathways and upstream regulators associated with the genetic changes reflective of eNOS functionality and dietary condition. The multigroup comparison across all four conditions identifies genes significantly associated with ‘Pathogen Induced Cytokine Storm’ and ‘Neuroinflammation’ signaling pathways (**Figure 5B**). Other canonical pathways include ‘Cardiac Hypertrophy’ and ‘PI3K/AKT’ signaling, likely reflecting the known effects of impaired eNOS function, and accordingly PI3K/Akt dysfunction, on cardiomyocyte signaling^24^. These genetic changes also yield a signature where cytokines such as IFNγ and TNF, and transcription factors, namely STAT3, were identified as upstream regulators (**Figure 5C**). The enhanced lesion formation in eNOS S1176A mice upon a Western Diet challenge underscores the importance of eNOS phosphorylation status in vascular protection, where eNOS activation state together with a Western Diet challenge leads to unique aortic genetic signatures.

**Figure 5.**
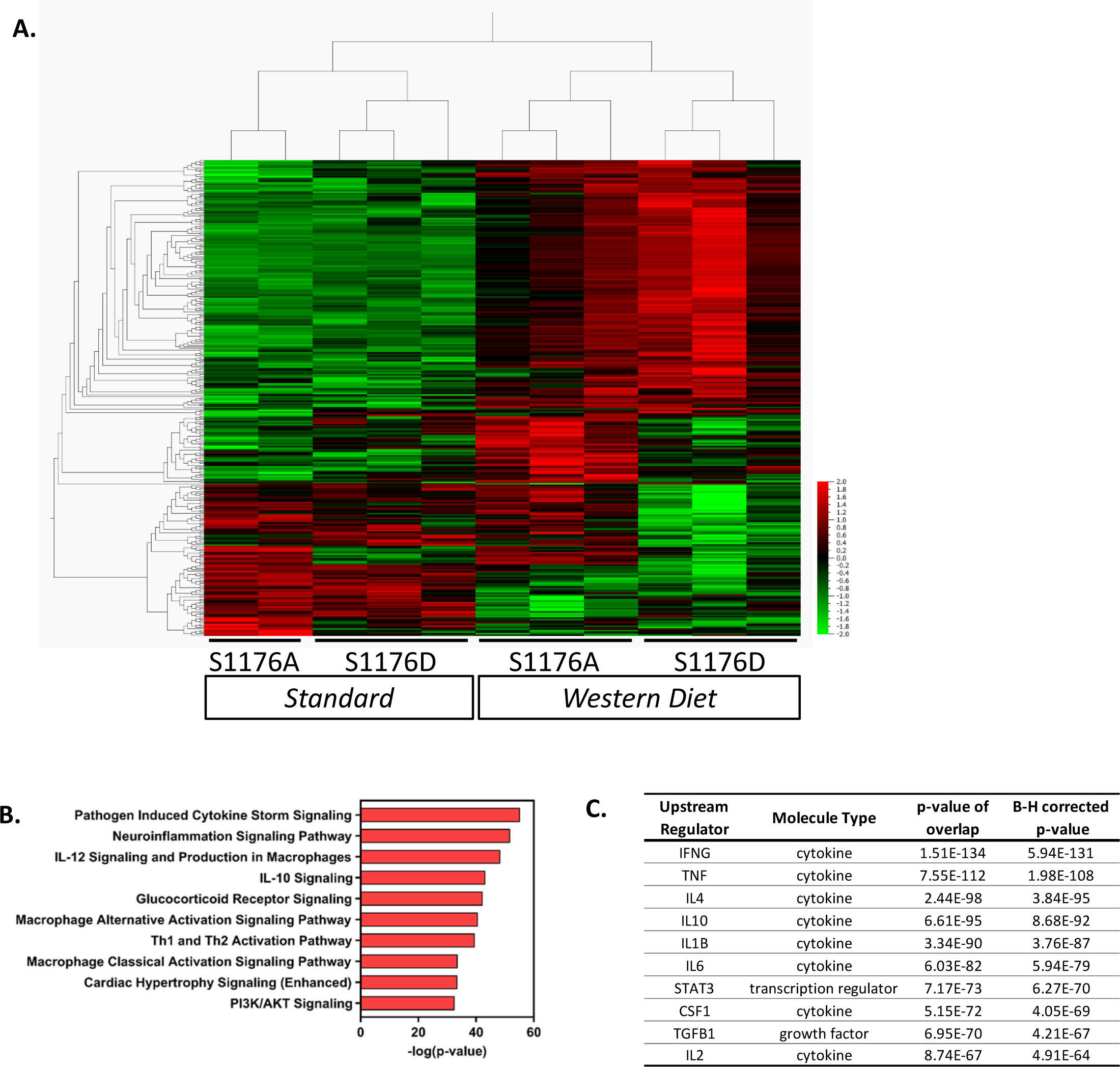
Gene expression analyses of whole aorta before and after 4-week Western Diet feeding. Gene expression profiles were obtained using the NanoString Immune Profile platform on whole aorta samples. **(A)** Heatmap of relative expression data illustrating unsupervised hierarchical gene clustering between comparison groups using multigroup comparison. Horizontal Columns: individual genes; Vertical Columns: individual mouse aorta samples. WD: Western Diet (p<0.05, 305 genes). **(B)** Pathway analyses identified several canonical pathways and **(C)** upstream regulators.

### Gene expression profile comparisons identify a subset of genes influenced by eNOS phosphorylation status rather than the environmental diet type

Previous studies using genetic epistasis approaches substantiated eNOS S1176 as the canonical phosphorylation site for the Akt-directed protective effects on vascular function *in vivo*. Mice harboring the endogenous eNOS S1176A and S1176D point-mutations were previously shown to express similar levels of eNOS protein, yet exhibit decreased and increased NO production, respectively^8^. We further analyzed the dataset to identify genetic changes associated with only eNOS phosphorylation status and unaffected by the given diet type. Two group analyses were performed to directly compare the eNOS mutant groups under conditions of either Standard or Western Diet feeding (p≤0.05). Under Standard diet conditions, we identify 82 differentially expressed genes in eNOS S1176A compared to eNOS S1176D mice (**Figure 6A**, 36 upregulated, 46 downregulated). Similar analysis focused on Western Diet fed conditions yield 118 genes in eNOS S1176A compared to S1176D mice (104 upregulated, 14 downregulated). Venn diagram analysis indicates a common set of 16 genes that are highly influenced by eNOS phosphorylation status (**Figure 6A**). Expression-based gene clustering indeed reveals these common genes to cluster with eNOS mutant status rather than diet type, as illustrated by the heatmap via sample clustering (**Figure 6B**). Further visualization of these shared genes using Box plot methods and Tukey analyses show significant changes associated with eNOS mutant status rather than diet type (**Supp. Fig. 5**). Interestingly, pathway analyses detect several canonical pathways associated with these transcriptional changes, including ‘Hepatic Fibrosis’, and ‘Rheumatoid Arthritis’ (**Figure 6C**). In accordance with eNOS activity as a protective mechanism, these pathways have previously been shown to associate with impaired eNOS function^25,26^. Given the significant impact of eNOS phosphorylation status on blood pressure regulation, we would predict these common genes to yield upstream regulators implicated in blood pressure control. As such, IPA analyses identifies ‘PRL’ and ‘NFAT5’ as the top predicted upstream regulators (**Fig 6D**). Previous studies have implicated high prolactin (PRL) levels with consequent hypertension^27^. Similarly, as a master transcriptional regulator responsive to hypertonicity, NFAT5 also plays a critical role in salt-dependent hypertension^28^. We therefore find that eNOS phosphorylation status, regardless of diet type, aligns with other mechanisms of blood pressure regulation.

**Figure 6.**
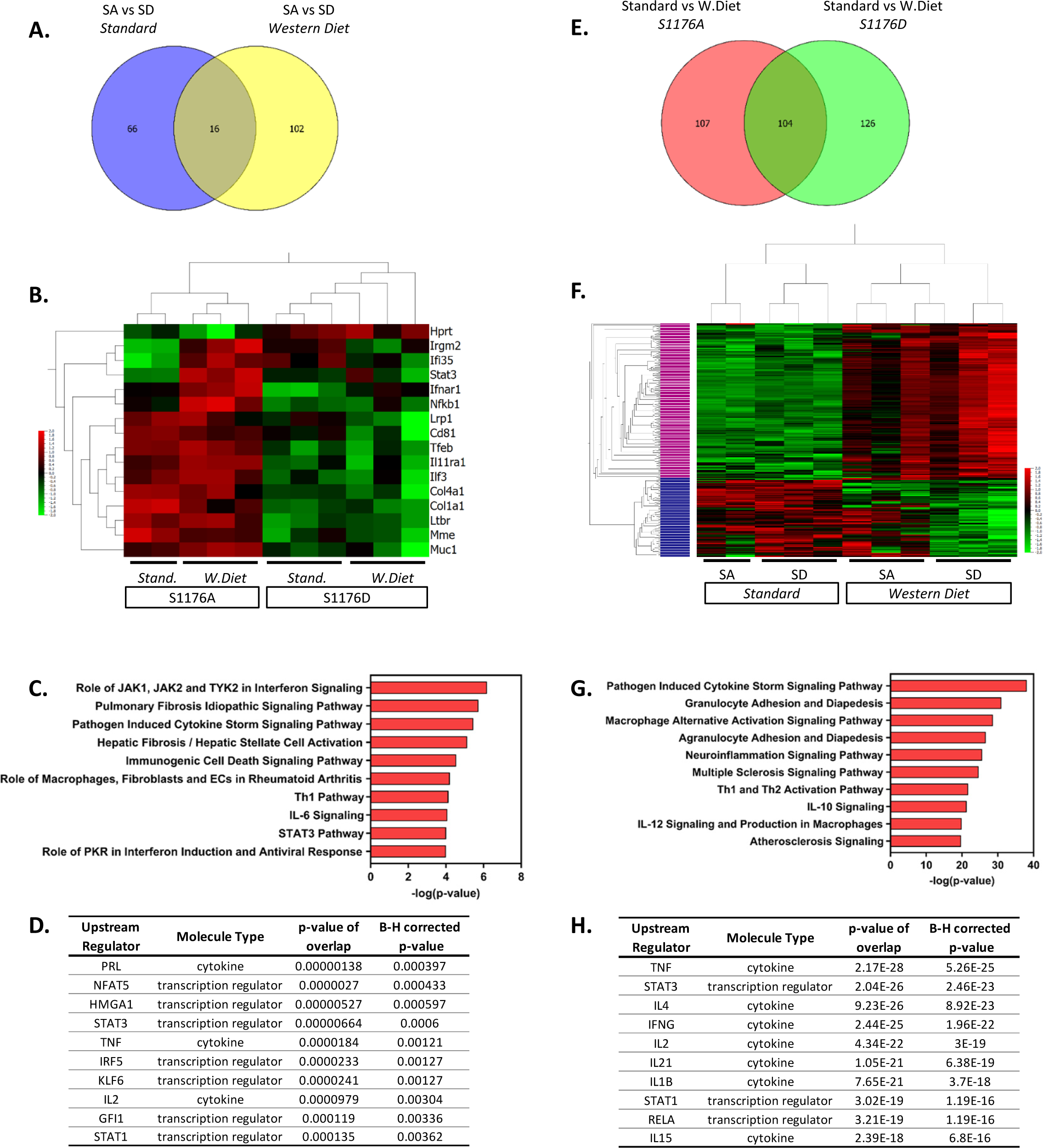
Venn diagram analysis of differentially expressed genes. Two-group comparisons were performed to allow for Venn diagram comparisons. **(A)** Venn diagram showing the number of differentially expressed genes between the two genetic groups (S1176A; S1176D) before and after Western Diet feeding. **(B)** Heatmap of the common genes driven by eNOS mutant across all sample groups. **(C)** Pathway analyses and **(D)** predicted upstream regulators of the common gene list from (A). **(E)** Venn diagram showing the number of differentially expressed genes between the two diet conditions (Standard, Western Diet) in the eNOS S1176A and S1176D group. **(F)** Heatmap of the common genes driven by diet type across all sample groups. **(G)** Pathway analyses of the upper cluster in heatmap from E (denoted in pink). **(H)** Predicted upstream regulators of commons genes from (E). Horizontal Columns: individual genes; Vertical Columns: individual mouse aorta samples.

### Gene expression profile comparison identifies a subset of genes influenced by diet type rather than eNOS phosphorylation status

eNOS-derived NO is a significant regulator of blood pressure, where loss of NO leads to hypertension, a well-known risk factor for atherosclerosis. Previous studies using pharmacological vasodilators in eNOS-deficient mice, however, did not mitigate atherosclerotic lesion development^29^. These findings indicate that hypertension does not fully account for the accelerated atherosclerosis in eNOS-deficient mice, suggesting alternative mechanisms of eNOS-mediated vascular protection. Our genetic mice provide a unique opportunity to identify eNOS-mediated protective mechanisms independent of blood pressure. We reasoned that blood pressure independent pathways would emerge by identifying differentially expressed genes influenced solely by diet type and unaffected by eNOS enzymatic status. We therefore performed two group comparisons to identify genes largely driven by environmental diet rather than eNOS phosphorylation status (p≤0.05). Analysis of only the phospho-impaired eNOS S1176A mice yielded 211 differentially expressed genes in Western diet versus Standard diet fed conditions (161 upregulated, 50 downregulated) (**Figure 6E**). Similar analysis of the eNOS S1176D mice identified 230 differentially expressed genes in Western diet versus Standard diet conditions (154 upregulated, 76 downregulated). Venn diagram analysis indicates a common set of 104 genes that are largely influenced by diet type rather than eNOS mutant form; these genes are differentially regulated by Western diet and not eNOS phosphorylation status (**Figure 6E**). Indeed, expression-based clustering of these common genes resulted in sample clustering into the two major diet groups (**Figure 6F**). Moreover, hierarchical clustering of these common genes reveals a major gene cluster, where expression levels are clearly upregulated under Western diet conditions in both eNOS mutant groups (**Figure 6F**, upper cluster denoted in pink, 155 genes). This cluster of genes may therefore elucidate pathways of atherosclerosis-related vascular inflammation that occur independently of eNOS activity. This upper gene cluster was subject to pathway analyses, where identified pathways include ‘Granulocyte/Agranulocyte Adhesion and Diapedesis’, ‘Multiple Sclerosis’, and ‘Atherosclerosis’ signaling pathways (**Figure 6G**). Along with previously identified regulators, IPA also determined IL-1β as an upstream activator unique to the list of genes influenced solely by diet type (**Figure 6H**). IL-1β is a well-known local and systemic contributor to cardiovascular inflammation, where the recent CANTOS clinical trial outcomes affirm IL-1β as an atherosclerosis-relevant inflammatory target^30^. The identification of ‘Atherosclerosis’ and ‘IL-1β’ as an upstream regulator for this shared cohort of genes thereby confirms the presence of an atherosclerotic gene signature.

### Gene expression analyses identify a unique set of differentially expressed genes in eNOS S1176A; Akt1^-/-^;ApoE^-/-^ mice on a Western Diet

Decades of work establish eNOS-derived NO as a cardioprotective agent both *in vitro* and *in vivo*^31–33^. As such, inhibition of NO in endothelial cells leads to increased expression of inflammatory adhesion molecules, where eNOS^-/-^ mice exhibit increased endothelial-leukocyte adhesion^32,34^. It is therefore likely that impaired eNOS phosphorylation and the consequent reduction in NO similarly leads to increased baseline vascular inflammation; eNOS S1176A mice may hence be primed for atherosclerosis to yield a unique gene signature upon a Western Diet challenge. Indeed, using Venn Diagram analyses of our two group comparisons, we identify a subset of 23 genes uniquely upregulated due to the combined effect of both ‘loss-of-function’ eNOS S1176A mutation and a Western Diet feeding period of 4 weeks (**Figure 7A**, asterisk). Indeed, a heatmap of these unique genes across the four sample groups clearly shows that the majority of these genes are upregulated only in the eNOS S1176A group when placed on a Western Diet, as all other groups do not indicate gene upregulation (**Figure 7B**). Visualization of these genes using Box plot methods and Tukey analyses show significant changes unique to the eNOS S1176A with a Western Diet challenge (**Supp. Fig. 6**). Further analyses of the identified 23 genes using IPA yielded several canonical pathways, including ‘April Mediated Signaling’, ‘B Cell Activating Factor’, ‘Complement System’, and ‘Regulation of the EMT by Growth Factors’ (**Figure 7C**). The cluster of enriched genes is predicted to be the result of several key inflammatory regulators, namely through the effects of the TNF, IL3, and IFNγ cytokines (**Figure 7D**). Pathways analyses also implicate the STAT and NFkB/RelA transcription factors, and is likely engaged for the downstream transcriptional effects of heightened cytokines, such as TNF, IL3, and IFNγ^35,36^. Accordingly, immunofluorescence staining of brachiocephalic lesions from eNOS S1176A mice display significantly increased Stat3 levels when compared to eNOS S1176D mice after a 12wk Western diet feeding period (**Figure 7E & 7F**). Immunofluorescence staining of RelA in eNOS S1176A mice also shows significant increases in expression when compared to eNOS S1176D mice (**Figure 7G & 7H**), thereby validating our gene expression analyses. The enhanced lesion formation in eNOS S1176A mice upon Western Diet challenge underscores the importance of eNOS phosphorylation status and function to maintain an athero-protective environment.

**Figure 7.**
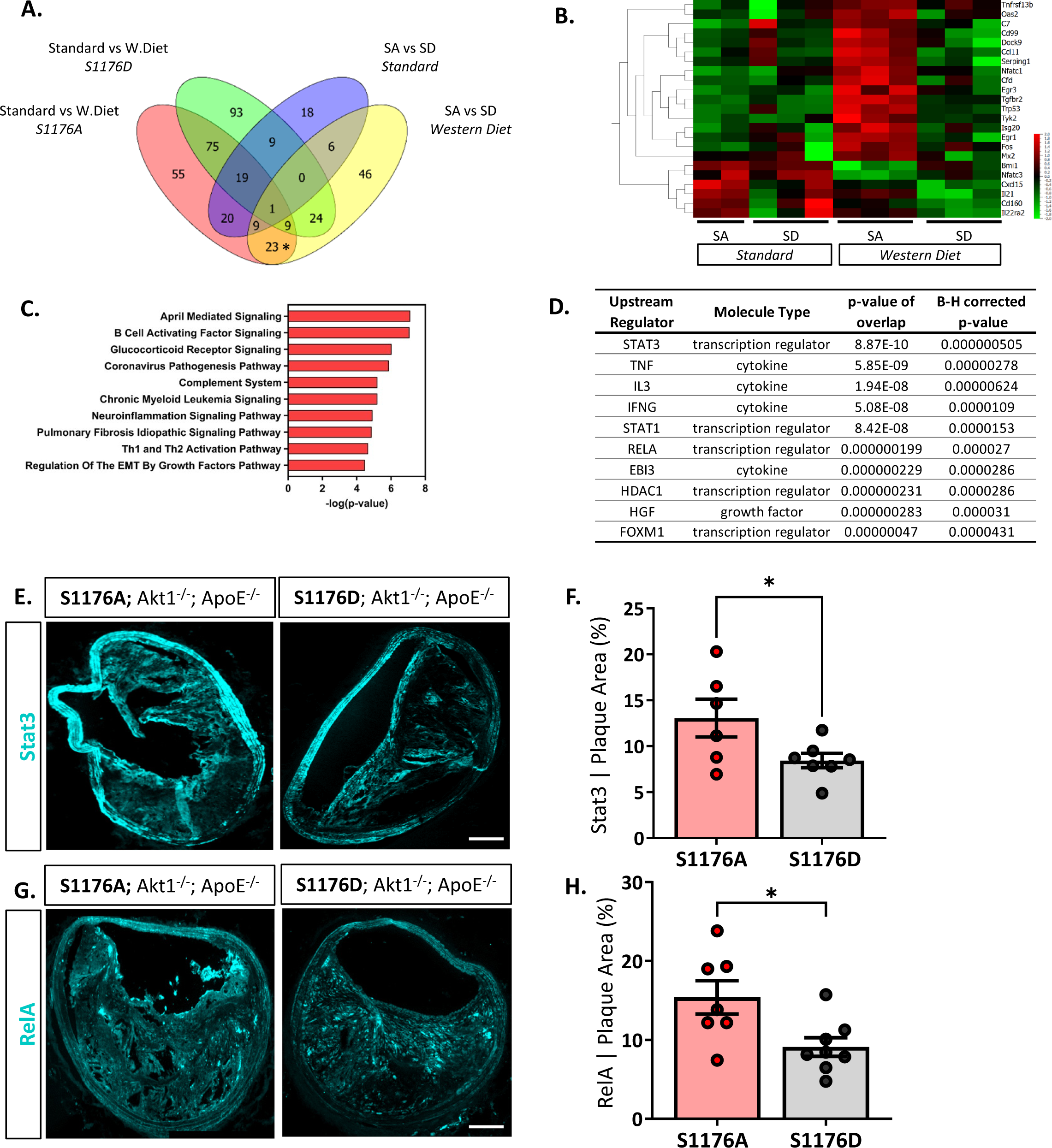
A‘loss-of-function’ mutation together with 4-week Western Diet feeding leads to a combinatorial increase in a select cohort of interferon-associated genes. **(A)** Venn diagram comparisons identify a subset of genes unique to combinatorial effects of impaired eNOS phosphorylation and a Western diet challenge (* on Venn). **(B)** Identified unique genes shown as a heatmap across all samples groups. **(C)** Pathway analyses identifies several canonical pathways and **(D)** predicted upstream regulators. Independent validation verifies an increased expression of **(E)** Stat3 and **(G)** RelA expression in eNOS S1176A compared to S1176D mice after a 12wk Western Diet challenge. Quantified in **(F&H)**. n=6 to 8 per group *, *p<0.05*

### Tumor Necrosis Factor (TNF) and Interferon-gamma (IFNγ)-driven pathways are elevated in atherosclerotic ‘loss-of-function’ eNOS S1176A mutant mice

Pathway analyses repeatedly identify TNF and IFNγ as two inflammatory and immune-activation related cytokines as upstream activators of the differentially expressed gene signature unique to the combinatorial effects of impaired eNOS (eNOS S1176A) and Western Diet challenge (**Figures 5C & 7D**). We therefore generated the predicted pathways via IPA and superimposed the 23 unique genes identified using Venn Diagram analyses to visualize the network driven by these two cytokines (**Figure 8**). As shown, many of these unique genes are predicted to be activated by TNF or IFNγ, suggesting that T-cell activation may drive the increased atherogenesis in the eNOS S1176A mice. More importantly, this study clarifies the existing discrepancy in the field surrounding eNOS function in atherosclerosis. By using elegant genetic epistasis approaches for i*n vivo* investigation of the Akt-eNOS axis in atherogenesis, our results affirm that eNOS S1176 phosphorylation is necessary to maintain an atheroprotective environment. Furthermore, our findings indicate that augmenting eNOS S1176 phosphorylation can overcome the well-known vascular consequences of impaired Akt1 expression, a kinase critical for endothelial protection.

**Figure 8:**
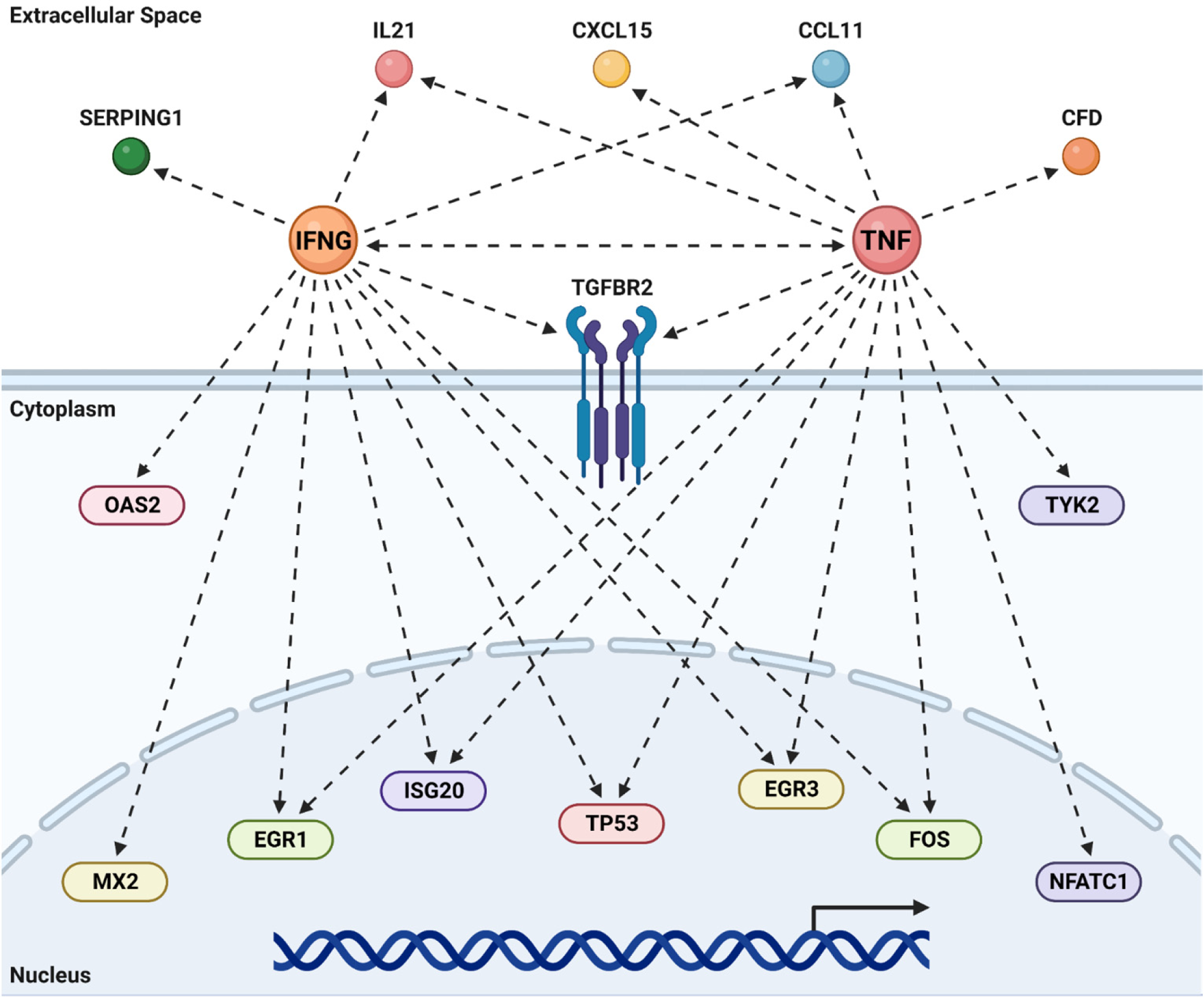
TNF & IFNγ promote atherogenesis in the absence of bioavailable NO. Predicted upstream regulator pathways derived from IPA were studied to derive the networks shown above. The genes unique to the combination of impaired eNOS phosphorylation (S1176A) and a Western Diet (Fig 6B) were super-imposed on these canonical pathways to look for agreement between real data and prediction. Most of the common genes are predicted by IPA to be under the regulation of either an activated TNF or IFNγ. Recreated in Biorender.

## DISCUSSION

The present study provides conclusive evidence through genetic epistasis approaches the importance of the Akt-eNOS kinase-substrate relationship for proper eNOS enzymatic function and cardiovascular benefit. Analysis at anatomical regions predisposed for atherosclerotic plaque formation indicate that mice harboring an unphosphorylatable (S1176A) eNOS point-mutation at the S1176 site results in an unfavorable lipoprotein profile, increased lipid deposition, cellular apoptosis, and increased inflammation, where pathway analyses identify preferential engagement of the TNF and IFNγ pathways. These deleterious effects are drastically reduced in the phospho-mimetic eNOS S1176D mice, suggesting a direct and significant correlation between eNOS enzymatic activity and atherogenic capacity. Through genetic modification of the eNOS S1176 activation site, we show that Akt1-mediated eNOS phosphorylation is critical for potentiating the vasculoprotective effects of endothelial Akt-eNOS signaling when challenged with dietary conditions that promote atherosclerosis.

Hypercholesterolemia promotes impaired endothelial-dependent vasorelaxation, where decreased NO bioavailability is an early characteristic of atherosclerosis^37^. While supplementation with eNOS substrates and cofactors enhance eNOS-derived NO production, an imbalance in eNOS enzymatic activity and cofactor levels can lead to eNOS uncoupling and detrimental effects^38,39^. Furthermore, mice overexpressing eNOS paradoxically show increased atherosclerotic lesion formation, emphasizing the complexity of eNOS regulation^6^. We therefore addressed the importance of Akt-mediated eNOS S1176 phosphorylation in limiting atherogenesis through site-specific modulation of eNOS enzymatic activity rather than protein levels. Our genetic comparison definitively shows that sustained eNOS S1176 phosphorylation is sufficient to mitigate atherosclerotic lesion formation, even in the absence of Akt1 expression, a critical cardioprotective kinase^40,41^. The decreased atherosclerotic plaque formation observed in eNOS S1176D mice reinforces eNOS S1176 phosphorylation as a therapeutic target for ameliorating vascular pathologies associated with endothelial dysfunction.

Our study provides the first assessment into the importance of the Akt1-eNOS activation cascade using genetic approaches. Despite the strengths of this study, there are several limitations. First, we acknowledge the omission of a eNOS^+/+^;Akt1^-/-^;ApoE^-/-^ double knockout (DKO) comparison group. The eNOS mutant mice (SA and SD) were bred to the identical eNOS^+/+^;Akt1^-/-^;ApoE^-/-^ DKO background from our previous reports, where we detail worsened atherosclerosis^13^. While we recognize the lack of a comparison group herein, our conclusion that eNOS activation decreases atherosclerosis remains the same. More importantly, atherosclerosis indices for the eNOS ‘constitutively active (S1176D; Akt1^-/-^; ApoE^-/-^)’ group herein are all less than what we previously reported for the eNOS^+/+^;Akt1^-/-^;ApoE^-/-^ DKO comparison group^13^. Secondly, tissue immunofluorescence staining did not include peptide blocking negative controls. However, the primary antibodies used for these studies were validated in previous publications and obtained through reputable commercial vendors (CD68-PMID:19002167; VCAM1-PMID:7530279; ITGA5-PMID:8491181; Stat3-PMID:29379096; RelA-PMID:32291445; Mac2-PMID:26948424). Lastly, it is possible that the atheroprotective effects of eNOS S1176D ‘gain-of-function’ are driven by both increased NO bioavailability and reduced plasma lipid levels. Unlike eNOS S1176A ‘loss-of-function’ mice, eNOS S1176D mice do not display an increase in plasma triglycerides with a Western Diet feed. Moreover, plasma triglyceride and cholesterol are significantly lower in eNOS S1176D mice, indicating that enhanced eNOS activity results in a favorable lipoprotein profile (**Figure 1C&1D),** similar to previous reports^14,42^. Despite these limitations, we find that a single mutation at eNOS S1176 yields divergent effects on atherosclerotic plaque formation.

Hypertension certainly plays a role in the development of atherosclerosis and related cardiovascular complications. Mice lacking both eNOS and ApoE display increased blood pressure when compared to ApoE-null mice^25^, suggesting that accelerated atherosclerosis may be due to systemic hypertension. However, treatment of eNOS/ApoE DKO with hydralazine, an ACE-independent vasodilator, resulted in lowered blood pressure with no effect on atherosclerotic lesion areas^29^. Furthermore, ApoE KO mice treated with levels of L-NAME low enough to maintain normotension also exhibit endothelial dysfunction and increased atherosclerotic lesions^43^. These studies suggest that NO deficiency promotes atherosclerosis through mechanisms independent of blood pressure regulation, such as the recruitment of inflammatory cells. Global deletion of Akt1 does not affect blood pressure^44^, likely due to the loss of Akt1 in other major cell types necessary for vascular tone (*e.g.* smooth muscle cells). The use of a global Akt1-null background together with the endogenous mutation of eNOS S1176 serves as a unique model, where we herein identify a genetic signature that begins to elucidate the blood pressure-independent consequences of eNOS deficiency. Furthermore, our genetic analyses identify several genes that are differentially affected by diet rather than eNOS activation status (**Figure 6F**), and may therefore yield potential targets of inflammation that are not sensitive to the beneficial effects of eNOS-mediated NO.

Atherosclerosis is often regarded as a chronic response to arterial tissue inflammation engaging both innate and adaptive immunity^45^, where NO has been shown to regulate such processes^46^. Evidence implicating the adaptive immune system has grown considerably in the last couple decades, as hypercholesterolemia can elicit a heighted activation state of T-cells. Moreover, atherosclerotic plaques are primarily associated with CD4+ T-cells characterized by heightened IFNy production^45^. Hence, immune responses mediated by the adaptive immune system may be the dominant force to promote inflammation in mature atherosclerotic lesions. Our pathways analyses indicate that the eNOS S1176 phosphorylation status modulates T-cell activity, as eNOS enzymatic impairment together with a Western Diet challenge leads to a combinatorial increase in select genes that are predicted to be a result of increased TNF and IFNγ levels. Our gene expression data also identifies ‘Regulation of the EMT Transition by Growth Factor Pathways’ as a top canonical pathway implicated under conditions of impaired eNOS function and Western Diet feeding (**Fig. 7C**). Elegant lineage-tracing studies have recently demonstrated that endothelial-to-mesenchymal (endoMT) transitioning occurs during atherosclerosis, where inflammatory conditions (*i.e.* TNF + IFNy) were shown to further promote endoMT processes^47,48^. Expression of the phospho-impaired eNOS S1176A predisposes the large vasculature to an inflammatory state where exposure to a Western diet promotes the preferential activation of TNF- and interferon-related pathways that may contribute to the recently described occurrences of endoMT in atherogenesis. This will, however, also require further investigation.

In conclusion, our model system herein investigates the effect of eNOS S1176 point-mutation on an atherogenic Akt1-null global deletion background^13^, where the genetic loss of Akt1 expression therefore prevents alternative protective effects mediated through endothelial Akt signaling. While eNOS S1176-independent mechanisms of NO production exist, our results suggest that Akt-mediated eNOS S1176 phosphorylation remains to be the most physiologically relevant method of eNOS activation to limit the vascular consequences of a Western Diet challenge. This study therefore shows not only the importance of maintaining eNOS S1176 phosphorylation, but the relevance of intact vascular Akt signaling, further reinforcing the physiological importance of the Akt1-eNOS kinase-substrate relationship for endothelial and vascular health.

## Supporting information

SuppFig

## ACKNOWLEDGEMENTS

We thank Paul L. Huang for providing the mutant eNOS mice for these studies. This work was supported in part by grants from the National Institutes of Health (grants R35 HL139945 and P01 HL1070205 to W.C.S, R00 HL130581 to M.Y.L.).

## AUTHOR CONTRIBUTIONS

M.Y.L. generated and characterized the mouse phenotypes. M.Y.L. performed the gene expression analyses. T.D.N. performed IPA analyses and validation experiments. M.Y.L. designed the project and experimental approaches. M.Y.L wrote the manuscript, which was reviewed and edited by all authors.

## FIGURE LEGENDS

**Supp. Fig. 1. PCA plots of aorta samples. 3-dimensional PCA plot showing the samples clustering according to their groups.** The PCA is based on 305 genes that are most variable between the groups using a multigroup comparison, for which the p-value is ≤ 0.05.

**Supp. Fig. 2. Whole aorta samples post-12wk Western Diet feeding.** Oil Red O staining for the whole aorta reflects a 12-week Western Diet feeding period. Multiple experimental cohorts shown.

**Supp. Fig. 3. Increased lipid deposition and cellular apoptosis in phospho-impaired eNOS-S1176A mutant mice. (A)** Oil Red O staining of the brachiocephalic artery 12 weeks post-Western Diet challenge. Lesion size and ORO quantified in **(B)** and **(C). (D)** TUNEL staining of the aortic root 12-weeks post-Western Diet, quantified in **(E)**. *n=5-6 per group*

**Supp. Fig. 4. Increased expression of Mac2 in phospho-deficient eNOS-S1176A mutant mice. (A)** Immunostaining of the brachiocephalic artery shows a trending increase in (A) Mac2 expression within atherosclerotic lesions of eNOS S1176A mice. Quantified in **(B).** *n=6 to 8 per group*

**Supp. Fig. 5. Box plots of genes from Figure 5B.** Output of Venn diagram analyses identifying differentially expressed genes in eNOS S1176A versus S1176D mutant group, and common to both Standard and Western Diet fed conditions.

**Supp. Fig. 6. Box plots of genes from Figure 6B.** Output of Venn diagram analyses identifying differentially expressed genes unique to the combined impact of impaired eNOS phosphorylation (S1176A) and a Western Diet challenge.

## REFERENCES

1. Liu VWT, Huang PL. Cardiovascular roles of nitric oxide: A review of insights from nitric oxide synthase gene disrupted mice†. Cardiovasc Res. 2008;77(1):19–29. doi:10.1016/j.cardiores.2007.06.024

2. Huang PL. Mouse Models of Nitric Oxide Synthase Deficiency. J Am Soc Nephrol. Published online 2000:4.

3. Kuhlencordt PJ, Gyurko R, Han F, et al. Accelerated Atherosclerosis, Aortic Aneurysm Formation, and Ischemic Heart Disease in Apolipoprotein E/Endothelial Nitric Oxide Synthase Double-Knockout Mice. Circulation. 2001;104(4):448–454. doi:10.1161/hc2901.091399

4. Moroi M, Zhang L, Yasuda T, et al. Interaction of genetic deficiency of endothelial nitric oxide, gender, and pregnancy in vascular response to injury in mice. J Clin Invest. 1998;101(6):1225–1232. doi:10.1172/JCI1293

5. Ozaki M, Kawashima S, Yamashita T, et al. Overexpression of endothelial nitric oxide synthase accelerates atherosclerotic lesion formation in apoE-deficient mice. J Clin Invest. 2002;110(3):331–340. doi:10.1172/JCI0215215

6. Kawashima S, Yokoyama M. Dysfunction of Endothelial Nitric Oxide Synthase and Atherosclerosis. Arterioscler Thromb Vasc Biol. 2004;24(6):998–1005. doi:10.1161/01.ATV.0000125114.88079.96

7. Atochin DN, Huang PL. Endothelial nitric oxide synthase transgenic models of endothelial dysfunction. Pflüg Arch - Eur J Physiol. 2010;460(6):965–974. doi:10.1007/s00424-010-0867-4

8. Schleicher M, Yu J, Murata T, et al. The Akt1-eNOS Axis Illustrates the Specificity of Kinase-Substrate Relationships in Vivo. Sci Signal. 2009;2(82):ra41–ra41. doi:10.1126/scisignal.2000343

9. Garcia V, Sessa WC. Endothelial NOS: perspective and recent developments. Br J Pharmacol. 2019;176(2):189–196. doi:10.1111/bph.14522

10. Rotllan N, Chamorro-Jorganes A, Araldi E, et al. Hematopoietic Akt2 deficiency attenuates the progression of atherosclerosis. FASEB J. 2015;29(2):597–610. doi:10.1096/fj.14-262097

11. Arranz A, Doxaki C, Vergadi E, et al. Akt1 and Akt2 protein kinases differentially contribute to macrophage polarization. Proc Natl Acad Sci. 2012;109(24):9517–9522. doi:10.1073/pnas.1119038109

12. Kerr BA, Ma L, West XZ, et al. Interference with Akt Signaling Protects Against Myocardial Infarction and Death by Limiting the Consequences of Oxidative Stress. Sci Signal. 2013;6(287):ra67–ra67. doi:10.1126/scisignal.2003948

13. Fernández-Hernando C, Ackah E, Yu J, et al. Loss of Akt1 Leads to Severe Atherosclerosis and Occlusive Coronary Artery Disease. Cell Metab. 2007;6(6):446–457. doi:10.1016/j.cmet.2007.10.007

14. Kashiwagi S, Atochin DN, Li Q, et al. eNOS phosphorylation on serine 1176 affects insulin sensitivity and adiposity. Biochem Biophys Res Commun. 2013;431(2):284–290. doi:10.1016/j.bbrc.2012.12.110

15. Cho H, Thorvaldsen JL, Chu Q, Feng F, Birnbaum MJ. Akt1/PKBα Is Required for Normal Growth but Dispensable for Maintenance of Glucose Homeostasis in Mice. J Biol Chem. 2001;276(42):38349–38352. doi:10.1074/jbc.C100462200

16. Shesely EG, Maeda N, Kim HS, et al. Elevated blood pressures in mice lacking endothelial nitric oxide synthase. Proc Natl Acad Sci. 1996;93(23):13176–13181. doi:10.1073/pnas.93.23.13176

17. Luo Y, Zhu Y, Basang W, Wang X, Li C, Zhou X. Roles of Nitric Oxide in the Regulation of Reproduction: A Review. Front Endocrinol. 2021;12:752410. doi:10.3389/fendo.2021.752410

18. Pallares P, Garcia-Fernandez RA, Criado LM, et al. Disruption of the endothelial nitric oxide synthase gene affects ovulation, fertilization and early embryo survival in a knockout mouse model. REPRODUCTION. 2008;136(5):573–579. doi:10.1530/REP-08-0272

19. Chen H, Howatt DA, Franklin MK, et al. A mini-review on quantification of atherosclerosis in hypercholesterolemic mice. Glob Transl Med. 2022;1(1):1–6. doi:10.36922/gtm.v1i1.76

20. Seimon TA, Wang Y, Han S, et al. Macrophage deficiency of p38α MAPK promotes apoptosis and plaque necrosis in advanced atherosclerotic lesions in mice. J Clin Invest. Published online March 16, 2009:JCI37262. doi:10.1172/JCI37262

21. Al-Yafeai Z, Yurdagul A, Peretik JM, Alfaidi M, Murphy PA, Orr AW. Endothelial FN (Fibronectin) Deposition by α5β1 Integrins Drives Atherogenic Inflammation. Arterioscler Thromb Vasc Biol. 2018;38(11):2601–2614. doi:10.1161/ATVBAHA.118.311705

22. Zhu SN, Chen M, Jongstra-Bilen J, Cybulsky MI. GM-CSF regulates intimal cell proliferation in nascent atherosclerotic lesions. J Exp Med. 2009;206(10):2141–2149. doi:10.1084/jem.20090866

23. Yuqing Huo, C. L. Ramos, K. Ley. Rolling and adhesion of monocytes to early atherosclerotic lesions of apolipoprotein E-/-(apoE-/-) mice requires P-selectin, PSGL-1, /spl alpha//sub 4/ integrin and VCAM-1. In: Proceedings of the First Joint BMES/EMBS Conference. 1999 IEEE Engineering in Medicine and Biology 21st Annual Conference and the 1999 Annual Fall Meeting of the Biomedical Engineering Society (Cat. N. Vol 1. ; 1999:56 vol.1. doi:10.1109/IEMBS.1999.802082

24. Yang Xiao-Ping, Liu Yun-He, Shesely Edward G., Bulagannawar Manohar, Liu Fang, Carretero Oscar A. Endothelial Nitric Oxide Gene Knockout Mice. Hypertension. 1999;34(1):24–30. doi:10.1161/01.HYP.34.1.24

25. Knowles JW, Reddick RL, Jennette JC, Shesely EG, Smithies O, Maeda N. Enhanced atherosclerosis and kidney dysfunction in eNOS–/–Apoe–/– mice are ameliorated by enalapril treatment. J Clin Invest. 2000;105(4):451–458. doi:10.1172/JCI8376

26. Totoson P, Maguin-Gaté K, Prati C, Wendling D, Demougeot C. Mechanisms of endothelial dysfunction in rheumatoid arthritis: lessons from animal studies. Arthritis Res Ther. 2014;16(1):R22. doi:10.1186/ar4450

27. Chang AS, Grant R, Tomita H, Kim HS, Smithies O, Kakoki M. Prolactin alters blood pressure by modulating the activity of endothelial nitric oxide synthase. Proc Natl Acad Sci. 2016;113(44):12538–12543. doi:10.1073/pnas.1615051113

28. Hiramatsu A, Izumi Y, Eguchi K, et al. Salt-Sensitive Hypertension of the Renal Tubular Cell– Specific NFAT5 (Nuclear Factor of Activated T-Cells 5) Knockout Mice. Hypertension. 2021;78(5):1335–1346. doi:10.1161/HYPERTENSIONAHA.121.17435

29. Chen J, Kuhlencordt PJ, Astern J, Gyurko R, Huang PL. Hypertension Does Not Account for the Accelerated Atherosclerosis and Development of Aneurysms in Male Apolipoprotein E/Endothelial Nitric Oxide Synthase Double Knockout Mice. Circulation. 2001;104(20):2391–2394. doi:10.1161/hc4501.099729

30. Ridker PM, Everett BM, Thuren T, et al. Antiinflammatory Therapy with Canakinumab for Atherosclerotic Disease. N Engl J Med. 2017;377(12):1119–1131. doi:10.1056/NEJMoa1707914

31. Lloyd-Jones, M.D DM, Bloch, M.D KD. THE VASCULAR BIOLOGY OF NITRIC OXIDE AND ITS ROLE IN ATHEROGENESIS. Annu Rev Med. 1996;47(1):365–375. doi:10.1146/annurev.med.47.1.365

32. De Caterina R, Libby P, Peng HB, et al. Nitric oxide decreases cytokine-induced endothelial activation. Nitric oxide selectively reduces endothelial expression of adhesion molecules and proinflammatory cytokines. J Clin Invest. 1995;96(1):60–68. doi:10.1172/JCI118074

33. Qian H, Neplioueva V, Shetty GA, Channon KM, George SE. Nitric Oxide Synthase Gene Therapy Rapidly Reduces Adhesion Molecule Expression and Inflammatory Cell Infiltration in Carotid Arteries of Cholesterol-Fed Rabbits. Circulation. 1999;99(23):2979–2982. doi:10.1161/01.CIR.99.23.2979

34. Lefer DJ, Jones SP, Girod WG, et al. Leukocyte-endothelial cell interactions in nitric oxide synthase-deficient mice. Am J Physiol-Heart Circ Physiol. 1999;276(6):H1943–H1950. doi:10.1152/ajpheart.1999.276.6.H1943

35. Pfeffer LM. The Role of Nuclear Factor κB in the Interferon Response. J Interferon Cytokine Res. 2011;31(7):553–559. doi:10.1089/jir.2011.0028

36. Reddy EP, Korapati A, Chaturvedi P, Rane S. IL-3 signaling and the role of Src kinases, JAKs and STATs: a covert liaison unveiled. Oncogene. 2000;19(21):2532–2547. doi:10.1038/sj.onc.1203594

37. Steinberg Helmut O., Bayazeed Basel, Hook Ginger, Johnson Ann, Cronin Jessica, Baron Alain D. Endothelial Dysfunction Is Associated With Cholesterol Levels in the High Normal Range in Humans. Circulation. 1997;96(10):3287–3293. doi:10.1161/01.CIR.96.10.3287

38. Hattori Y, Hattori S, Wang X, Satoh H, Nakanishi N, Kasai K. Oral Administration of Tetrahydrobiopterin Slows the Progression of Atherosclerosis in Apolipoprotein E-Knockout Mice. Arterioscler Thromb Vasc Biol. 2007;27(4):865–870. doi:10.1161/01.ATV.0000258946.55438.0e

39. Cosentino F, Katusic ZS. Tetrahydrobiopterin and Dysfunction of Endothelial Nitric Oxide Synthase in Coronary Arteries. Circulation. 1995;91(1):139–144. doi:10.1161/01.CIR.91.1.139

40. Lee MY, Luciano AK, Ackah E, et al. Endothelial Akt1 mediates angiogenesis by phosphorylating multiple angiogenic substrates. Proc Natl Acad Sci. 2014;111(35):12865–12870. doi:10.1073/pnas.1408472111

41. Shiojima I, Walsh K. Role of Akt Signaling in Vascular Homeostasis and Angiogenesis. Circ Res. 2002;90(12):1243–1250. doi:10.1161/01.RES.0000022200.71892.9F

42. Van Haperen R, De Waard M, Van Deel E, et al. Reduction of Blood Pressure, Plasma Cholesterol, and Atherosclerosis by Elevated Endothelial Nitric Oxide. J Biol Chem. 2002;277(50):48803–48807. doi:10.1074/jbc.M209477200

43. Kauser K, da Cunha V, Fitch R, Mallari C, Rubanyi GM. Role of endogenous nitric oxide in progression of atherosclerosis in apolipoprotein E-deficient mice. Am J Physiol-Heart Circ Physiol. 2000;278(5):H1679–H1685. doi:10.1152/ajpheart.2000.278.5.H1679

44. Symons JD, McMillin SL, Riehle C, et al. Contribution of Insulin and Akt1 Signaling to Endothelial Nitric Oxide Synthase in the Regulation of Endothelial Function and Blood Pressure. Circ Res. 2009;104(9):1085–1094. doi:10.1161/CIRCRESAHA.108.189316

45. Wolf D, Ley K. Immunity and Inflammation in Atherosclerosis. Circ Res. 2019;124(2):315-327. doi:10.1161/CIRCRESAHA.118.313591

46. Bogdan C. Nitric oxide synthase in innate and adaptive immunity: an update. Trends Immunol. 2015;36(3):161–178. doi:10.1016/j.it.2015.01.003

47. Evrard SM, Lecce L, Michelis KC, et al. Endothelial to mesenchymal transition is common in atherosclerotic lesions and is associated with plaque instability. Nat Commun. 2016;7(1):11853. doi:10.1038/ncomms11853

48. Chen PY, Qin L, Baeyens N, et al. Endothelial-to-mesenchymal transition drives atherosclerosis progression. J Clin Invest. 2015;125(12):4514–4528. doi:10.1172/JCI82719

