## Supplementary material for "Endothelial Nitric Oxide Synthase (eNOS) S1176 phosphorylation status governs atherosclerotic lesion formation": SuppFig

***Supplemental Figure 1***

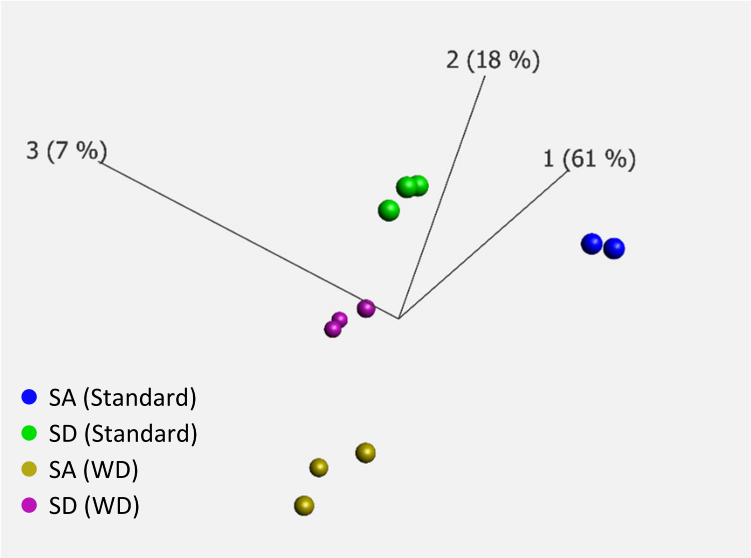


**Supp. Fig. 1. PCA plots of aorta samples.** 3-dimensional PCA plot showing the samples clustering according to their groups. The PCA is based on 305 genes that are most variable between the groups using a multigroup comparison, for which the p-value is ≤ 0.05.

#### *Supplemental Figure 2*

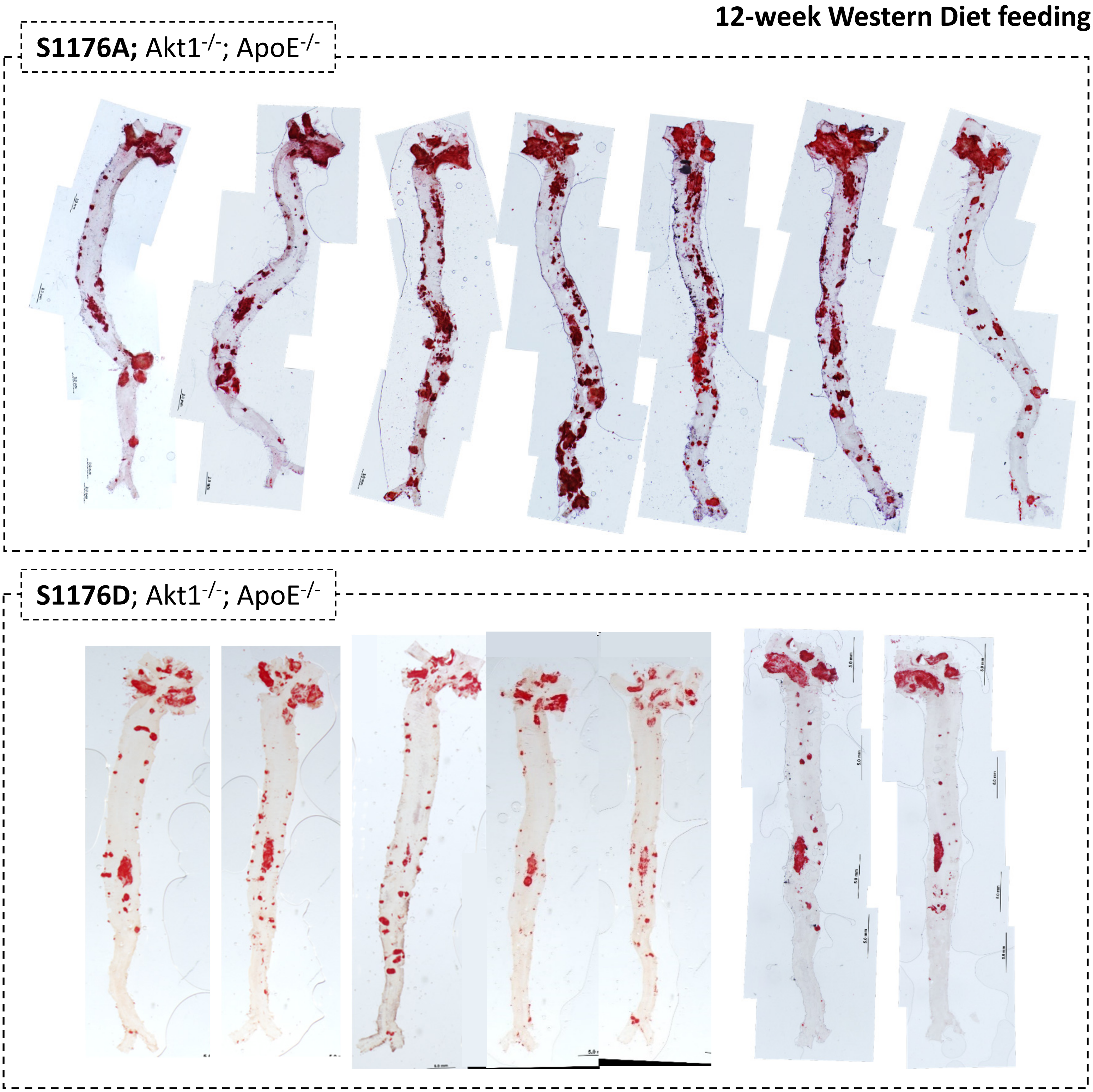


**Supp. Fig. 2. Whole aorta samples post-12wk Western Diet feeding.** Oil Red O staining for the whole aorta reflects a 12-week Western Diet feeding period. Multiple experimental cohorts shown.

### Supplemental Figure 3

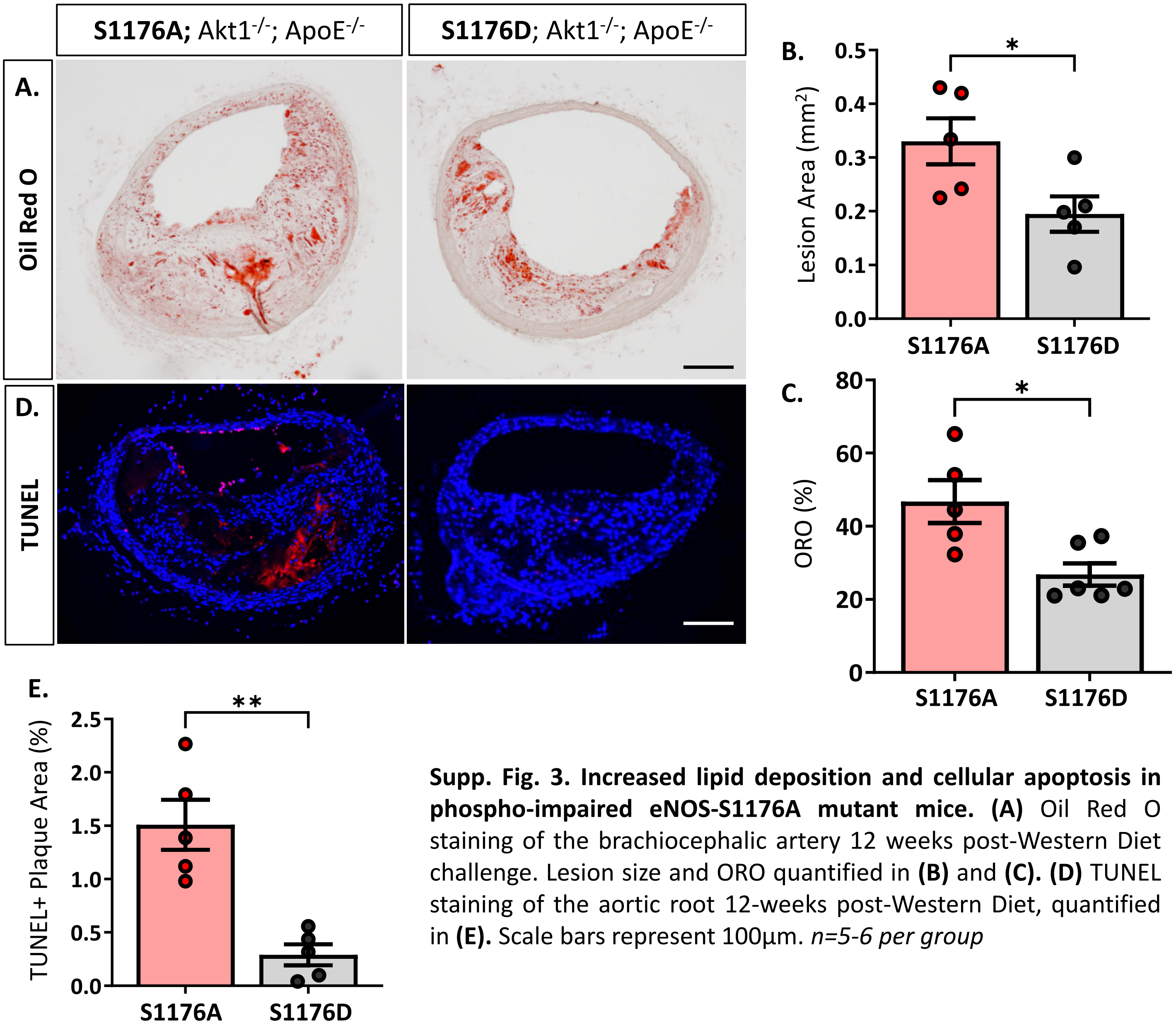


**Supp. Fig. 3. Increased lipid deposition and cellular apoptosis in phospho-impaired eNOS-S1176A mutant mice. (A)** Oil Red O staining of the brachiocephalic artery 12 weeks post-Western Diet challenge. Lesion size and ORO quantified in **(B)** and **(C). (D)** TUNEL staining of the aortic root 12-weeks post-Western Diet, quantified in **(E).** Scale bars represent 100µm. *n=5-6 per group*

### ***Supplemental Figure 4***

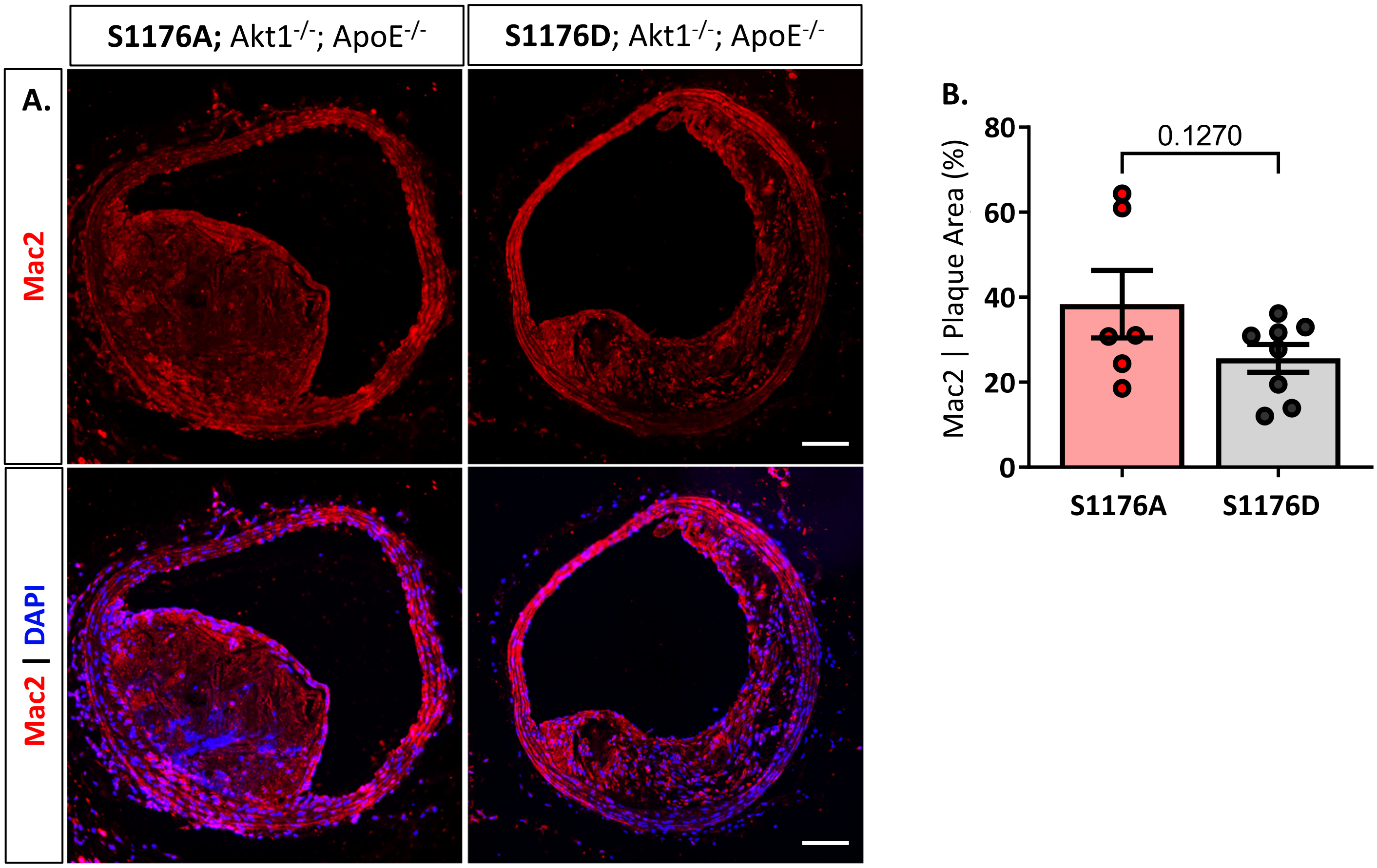


**Supp. Fig. 4. Increased expression of Mac2 in phospho-deficient eNOS-S1176A mutant mice. (A)** Immunostaining of the brachiocephalic artery shows a trending increase in **(A)** Mac2 expression within atherosclerotic lesions of eNOS S1176A mice. Scale bar represents 100µm. Quantified in **(B)**. *n=6-8 per group*

### ***Supplemental Figure 5***

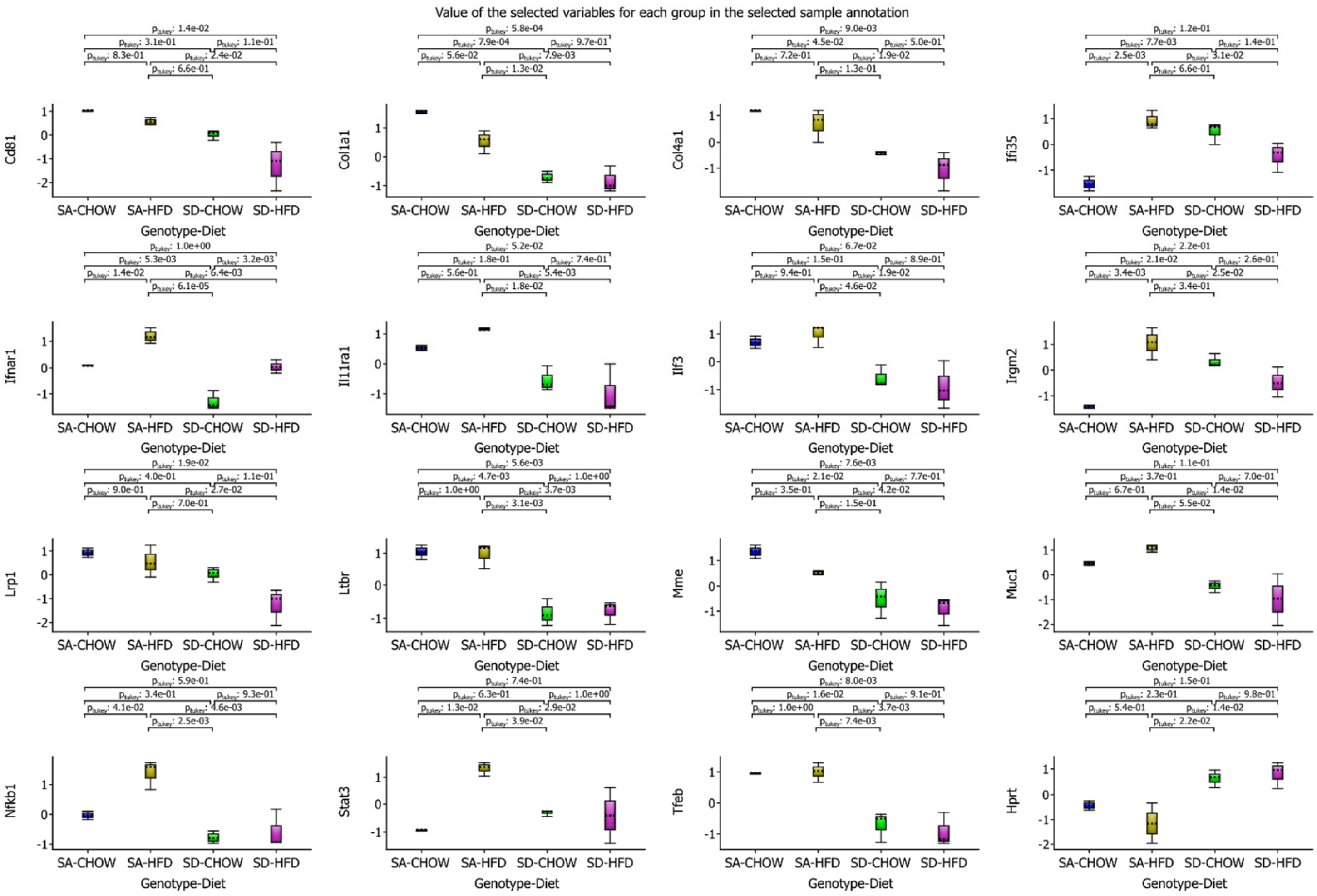


**Supp. Fig. 5. Box plots of genes from Figure 5B.** Output of Venn diagram analyses identifying differentially expressed genes in eNOS S1176A versus S1176D mutant group, and common to both Standard and Western Diet fed conditions.

### ***Supplemental Figure 6***

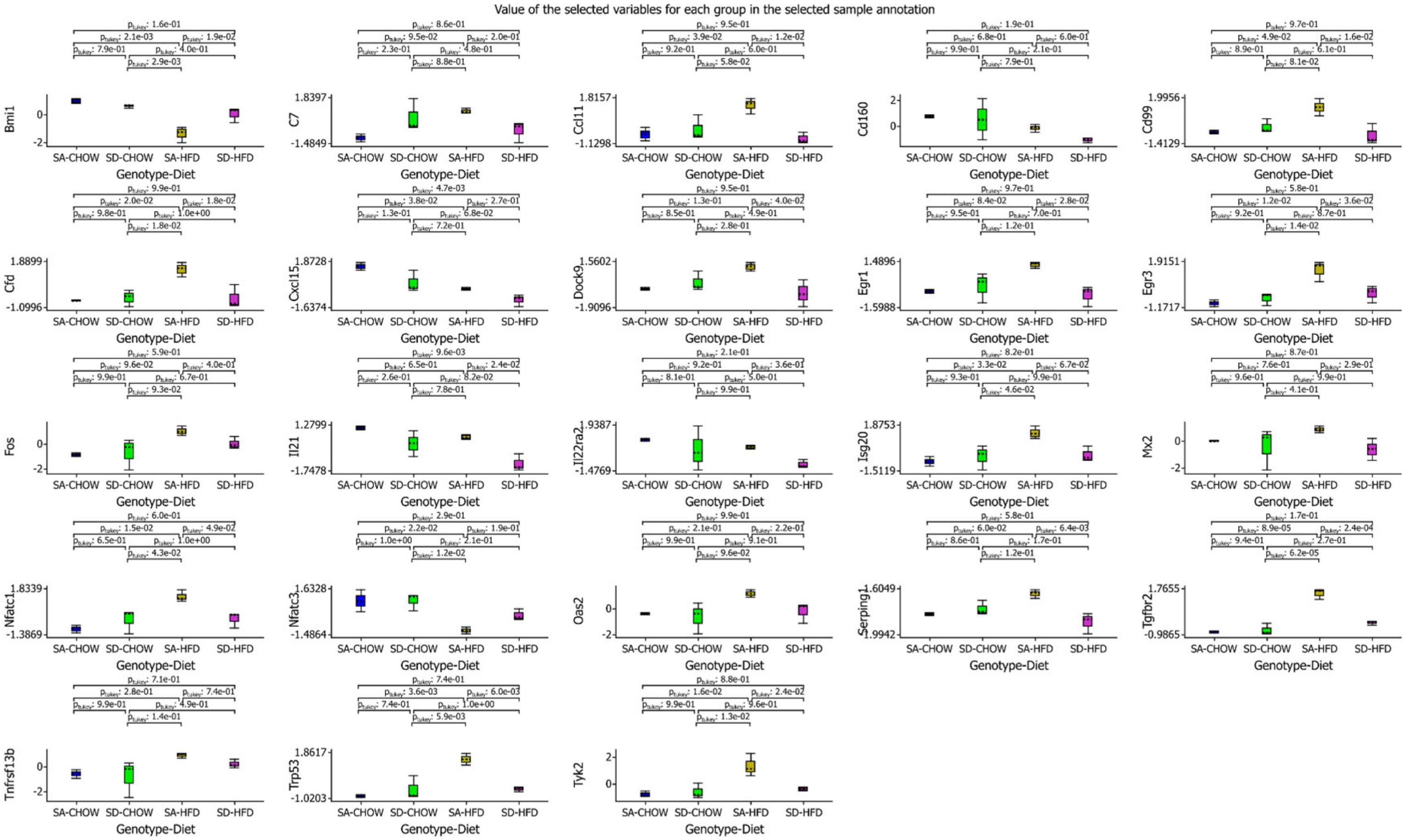


**Supp. Fig. 6. Box plots of genes from Figure 6B.** Output of Venn diagram analyses identifying differentially expressed genes unique to the combined impact of impaired eNOS phosphorylation (S1176A) and a Western Diet challenge.
